# SPLENDID incorporates continuous genetic ancestry in biobank-scale data to improve polygenic risk prediction across diverse populations

**DOI:** 10.1101/2024.10.14.618256

**Authors:** Tony Chen, Haoyu Zhang, Rahul Mazumder, Xihong Lin

## Abstract

Polygenic risk scores are widely used in disease risk stratification, but their accuracy varies across diverse populations. Recent methods large-scale leverage multi-ancestry data to improve accuracy in under-represented populations but require labelling individuals by ancestry for prediction. This poses challenges for practical use, as clinical practices are typically not based on ancestry. We propose SPLENDID, a novel penalized regression framework for diverse biobank-scale data. Our method utilizes ancestry principal component interactions to model genetic ancestry as a continuum within a single prediction model for all ancestries, eliminating the need for discrete labels. In extensive simulations and analyses of 9 traits from the All of Us Research Program (N=224,364) and UK Biobank (N=340,140), SPLENDID significantly outperformed existing methods in prediction accuracy and model sparsity. By directly incorporating continuous genetic ancestry in model training, SPLENDID stands as a valuable tool for robust risk prediction across diverse populations and fairer clinical implementation.

## Introduction

Polygenic risk scores (PRS) are emerging as a promising tool for stratifying disease risk and guiding personalized intervention decisions^1–3^. However, most PRS models have been developed using data primarily from European ancestry populations ^4–7^, which have limited accuracy in individuals from non-European ancestries^8^. These discrepancies arise due to differences in genetic factors like minor allele frequencies (MAF), linkage disequilibrium (LD), and genetic effect sizes across populations ^9,10^. To address this, new methods have emerged that incorporate summary statistics from genome-wide association studies (GWAS) across multiple ancestries^11^. These approaches range from Bayesian priors^12–14^ such as PRS-CSx that jointly model ancestry effects, clumping-based methods such as CT-SLEB^15^ that use two-way p-value and LD thresholding, and penalized regression such as in PROSPER^16^ that can encourage shared sparsity and effect sizes. These methods take advantage of large sample sizes and statistical power from European-based GWAS, while also integrating ancestry-specific effects and LD structure to improve PRS accuracy to under-represented target populations.

While these multi-ancestry PRS methods have shown substantial improvements in prediction accuracy over single-ancestry analysis, they all focus on a discrete notion of ancestry. However, human genetic ancestry is a continuous spectrum and treating it as such is critical for improving the accuracy of polygenic scores across diverse populations ^17^. Recent work has shown that the accuracy of PRS declines not only between distinct ancestries, but also over a continuum of genetic ancestry, even within a single ancestry group^18^. Moreover, since individuals may not belong to pre-specified ancestry categories and clinical practices are generally not tailored to specific ancestries, ancestry-specific PRS have limited practical utility. This underscores the need for a more nuanced approach that reflects the complex continuum of genetic diversity, improving the feasibility of PRS in clinical practice.

Recent efforts have begun to develop continuous ancestry PRS models using individual-level data from diverse cohorts. One such approach, GAUDI^19^, utilizes local ancestry inference^20^ to separate individuals’ genotypes into ancestry-specific segments and applies fused Lasso regression to model ancestry-specific genetic effects. Although promising, GAUDI is limited to analysis of two-way admixed ancestry (i.e. European and African) and a relatively small number of individuals and genetic variants. Another approach, the inclusive polygenic score with ancestry-specific refitting (iPGS+refit)^21^, uses Lasso regression to model shared genetic effects among all ancestries, then refits an additional Lasso model in each ancestry group with genetic interactions with ancestry principal components (PCs) to model ancestry-dependent effects. Although PCs are an effective continuous measure of genetic ancestry, the ancestry-specific refitting procedure produces distinct prediction models for each ancestry and still requires pre-classifying subjects into discrete ancestry groups. Thus, there remains a gap in PRS methodology for accurate PRS prediction within a single label-free model that can be applied to individuals of any ancestry.

To address these limitations, we propose SPLENDID (**SP**arse **L**inear **EN**semble**D I**nteraction model with **D**ifferential ancestry genetic effects), a polygenic risk score method that models the relationship between genetic variants and ancestry on a continuous scale. SPLENDID uses a group-L0L1 penalty framework^22^ and ensemble learning^5,23^ to capture both shared and ancestry-specific genetic effects, without the need for discrete ancestry labels. This approach enables a single, unified prediction model that maintains robust accuracy across diverse populations, making the use of PRS in clinical practice more feasible.

In large-scale simulations and analysis of nine quantitative traits from the All of Us Research Program (AoU, N=224,364) and UK Biobank (UKBB, N=340,140), SPLENDID achieved similar or significantly higher prediction accuracy than existing methods, while producing highly sparse and interpretable PRS. By providing a more inclusive and flexible framework for PRS, SPLENDID represents an effective step in advancing personalized risk prediction for individuals from all ancestries and improving its practical utility in clinical settings.

## Results

### Method overview

SPLENDID is a penalized regression-based PRS method that incorporates interactions between genetic variants and ancestry PCs to capture heterogeneous genetic effects by treating ancestry as a continuum rather than using discrete ancestry labels (**Figure 1, Methods**). We utilize a group-L0L1 penalized regression framework^22^ to perform sparse selection of main genetic effects and GxPC interactions using a combinatorial L0 penalty with a group-level L1 shrinkage for out-of-sample prediction. While such regression problems are computationally challenging in principle, we use specialized algorithms for efficient analysis of biobank-scale data. We then use an ensemble learning strategy to optimally combine several candidate models and further enhance prediction accuracy across diverse ancestries, while maintaining sparsity for an interpretable prediction model.

**Figure 1.**
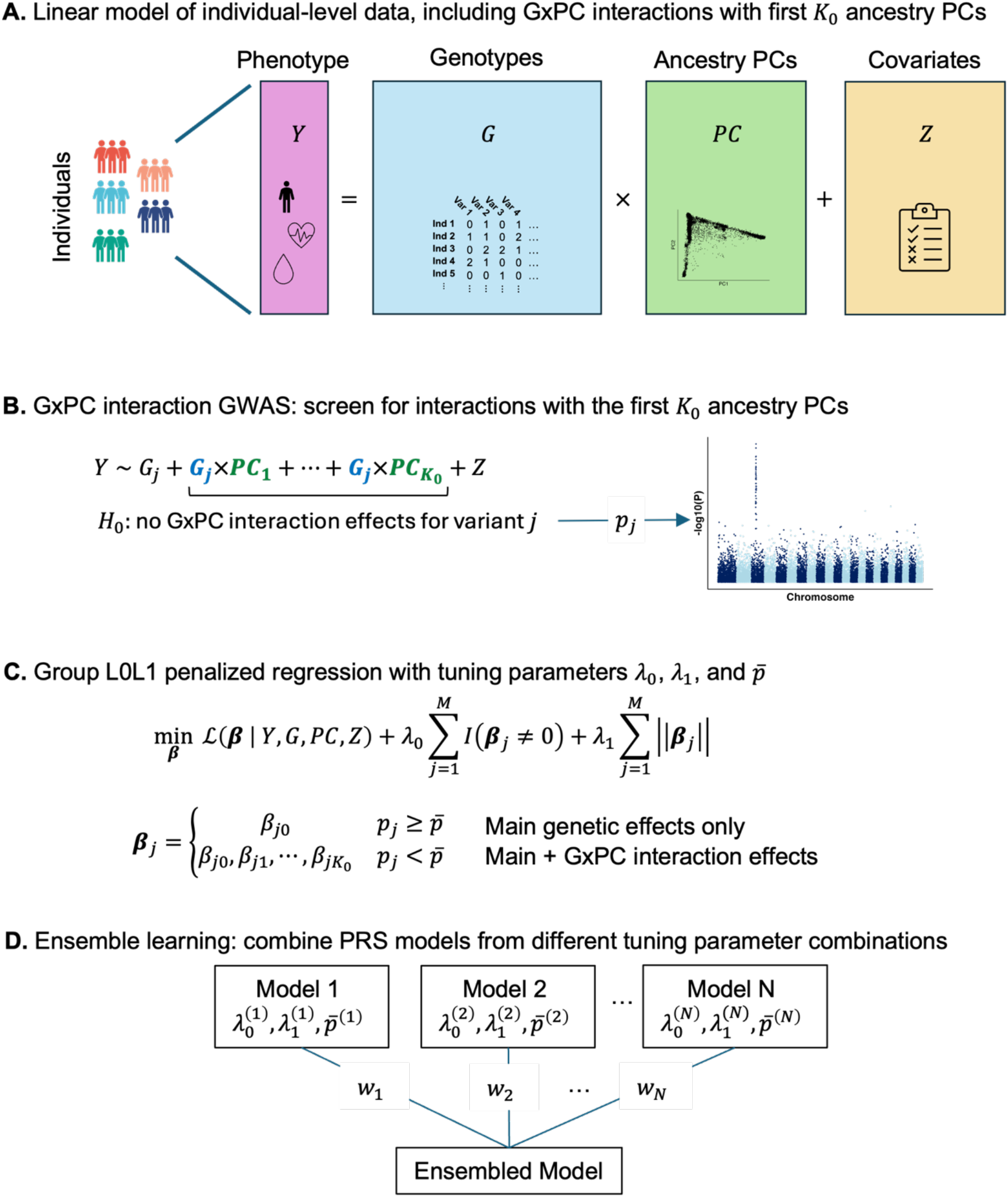
SPLENDID method overview. (A) Our method utilizes individual-level genotype and phenotype data in multi-ancestry biobanks, as well as ancestry PCs and other relevant covariates, such as age, sex, etc. In particular, we are interested in building a single multi-ancestry PRS model by modelling interactions between genetic variants ***G*** and ancestry PCs by treating ancestry as a continuum to detect effect heterogeneity across ancestries. (B) We first screen for interactions between each variant ***j*** and the first ***K***_**0**_ ancestry PCs using marginal GxPC interaction GWAS analysis to obtain p-values ***p***_***j***_ testing for GxPC interaction effects for different p-value thresholds 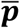. (C) We then use group-L0L1 penalized regression to fit the joint GxPC interaction model and select a sparse set of predictive variants and any GxPC interactions. Our approach takes in a grid of penalty parameters ***λ***_**0**_ and ***λ***_**1**_, as well as p-value threshold 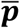 used to select GxPC interactions. (D) Finally, we use a sparse ensemble learning strategy to combining PRS from different combinations of ***λ***_**0**_, ***λ***_**1**_, and 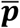 in an independent tuning dataset.

SPLENDID requires three independent datasets: training, tuning, and validation. In the training data, we model a continuous phenotype ***Y*** in terms of genetic variants ***G*** with effect sizes *B*_*ij*_ for individual *i* and variant *j*, the first *K* (e.g. 20) ancestry PCs, and covariates ***Z***.

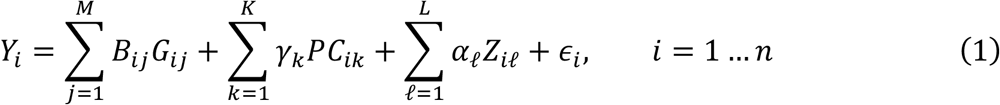

where the genetic effects *B*_*ij*_ may either be a constant value when effects are homogeneous or heterogeneous and vary across all ancestries as a function of genetic ancestry, which we can quantify using the first *K*_0_ (e.g. 5) ancestry PCs:

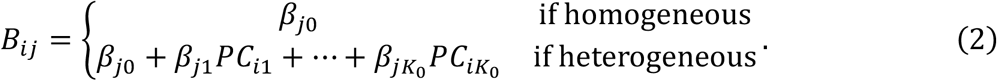

Using *H* to denote the set of variants with heterogeneous effects, Equation 1 then expands into the following GxPC interaction model:

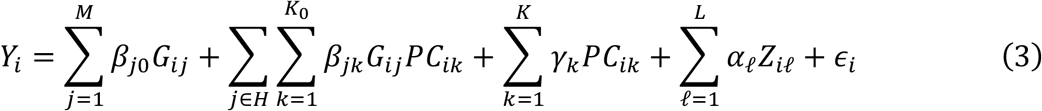

We first screen for relevant interactions to include in *H* using marginal GxPC interaction GWAS. We then run group-L0L1 penalized regression to sparsely using a several combinations of penalty parameters (*λ*_0_, *λ*_1_) and significance thresholds 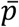 for the GxPC GWAS (**Methods**). The final prediction model is constructed by an ensemble learning strategy that optimizes overall prediction accuracy by flexibly weighting models from different combinations of *λ*_0_, *λ*_1_, and 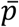 while also prioritizing sparsity for interpretability (**Methods**). This ensembled PRS model can then be applied to and evaluated in an independent validation dataset.

### Simulation studies

We evaluated the performance of SPLENDID using simulated individuals following realistic LD patterns from African (AFR), Admixed American (AMR), East Asian (EAS), European (EUR), and South Asian (SAS) ancestry based on 1000G and restricted to about 1.1 million variants from HapMap3^12,24^. We compared our method against multi-ancestry GWAS summary statistics based PRS methods PRS-CSx^12^, CT-SLEB^15^, and PROSPER^16^, as well as iPGS+refit^21^ which utilizes multi-ancestry individual-level data.

Simulated outcomes were generated with heritability 0.4 in each ancestry under various levels of trait polygenicity (0.05% to 1% causal) and patterns of effect heterogeneity (**Methods**). One heterogeneity pattern assumed all causal genetic effects followed a multivariate normal (MVN) distribution with “fixed” pairwise correlation *ρ* of 0.6 or 0.8 across the five ancestries. Another “mixed” pattern held half of the causal effects identical across all ancestries, with the other half following a MVN with pairwise correlation 0.4 or 0.6, corresponding to overall cross-ancestry genetic correlation of 0.7 or 0.8, respectively. For each simulation setting, we trained each method using 100,000 samples with varying proportions of non-European ancestry, tuned using 10,000 of European ancestry and 2,000 of each non-European ancestry (N=18,000), then validated in 10,000 samples of each ancestry (N=50,000).

When the proportions of causal genetic variants were low (0.05% and 0.1%), the average R^2^ for SPLENDID among all validation samples was 0.1% higher than PRS-CSx, 4% higher than CT-SLEB, 13.1% higher than PROSPER, and 12.5% higher than iPGS+refit across all ancestry proportion and heterogeneity pattern settings (**Figure 2, Supplementary Figure 1, Supplementary Figure 2, Supplementary Table 1**). In particular, under the mixed heterogeneity patterns, SPLENDID achieved substantially better prediction accuracy compared to all four alternative methods: 3.7% higher than PRS-CSx, 7.2% higher than CT-SLEB, 16.2% higher than PROSPER, and 12.3% higher than iPGS+refit. When causal effects followed a single MVN, PRS-CSx slightly outperformed SPLENDID, by about 3.4% on average.

**Figure 2.**
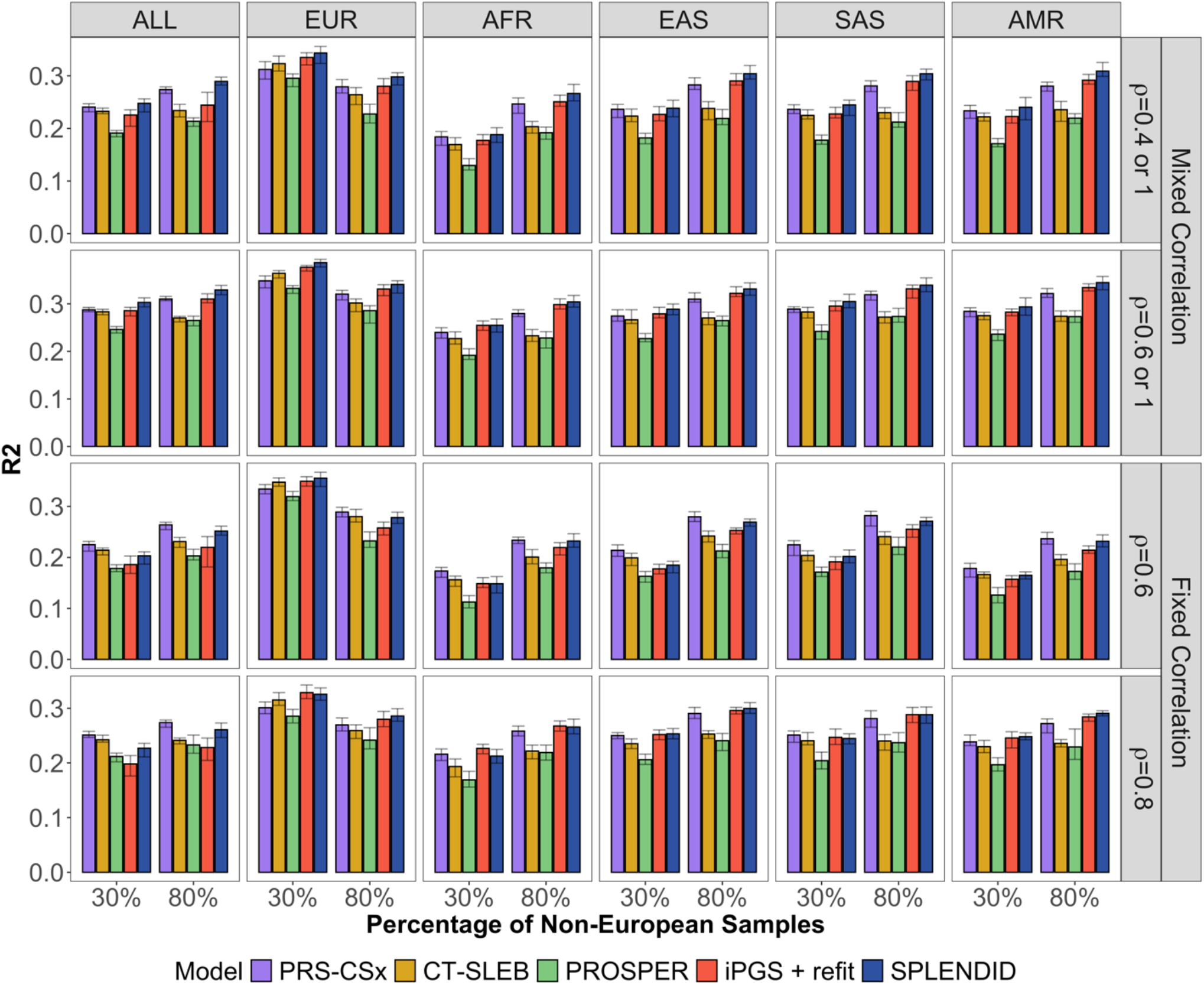
Prediction accuracy of PRS methods in simulated data. We compared SPLENDID with PRS-CSx, CT-SLEB, PROSPER, and iPGS+refit with a total training size of 100,000 samples, and 1,158 (0.1%) causal variants. We varied the genetic correlation structures and the proportion of non-European ancestry within the training dataset. Each method was tuned in an independent tuning dataset with 18,000 samples, and evaluated in a validation dataset with 50,000 samples (10,000 of each ancestry). Prediction R^2^ values were reported as the average over 10 simulation replicates for each setting, with error bars corresponding to the total range, among all 50,000 validation samples and stratified by each ancestry group.

In simulations with more polygenic outcomes (0.5% and 1% causal genetic variants), SPLENDID attained an average R^2^ among all validation samples that was 4.9% higher than PRS-CSx, 19.8% higher than CT-SLEB, and 20.2% higher than PROSPER, but 2.3% lower than iPGS+refit (**Supplementary Figure 3, Supplementary Figure 4**). In these scenarios, the magnitude of causal effects and any heterogeneity across ancestries were very small, making both main effects and interactions harder to detect. iPGS+refit tends to select a large number of variants, which may lend to better prediction in highly polygenic scenarios. Nonetheless, a key practical advantage for SPLENDID over the existing methods is that it provides a single prediction model that is applicable to all ancestries, rather than separate models for each ancestry, mimicking practical clinical settings.

PRS-CSx achieved higher accuracy than SPLENDID for fixed genetic correlation with 0.05% and 0.1% causal variants. However, when tuning PRS-CSx on the pooled tuning dataset to create a single PRS like SPLENDID, PRS-CSx had an average 10% lower R^2^ compared to the original ancestry-specific tuning, and 20% lower R^2^ compared to SPLENDID tuned in pooled ancestry. (**Supplementary Figure 5, Supplementary Table 2**). SPLENDID showed consistent accuracy between the two tuning approaches, with some slightly improvements from ancestry-specific tuning. This may be because ancestry-dependent effects are already captured during training, so that ensembling is less sensitive to the ancestry makeup in tuning. However, we still advocate for pooled ancestry tuning to provide a single, inclusive prediction model to improve its fair implementation in clinical settings.

In addition to improved and robust prediction accuracy across various underlying settings, SPLENDID recovered close to the true support size regardless of trait polygenicity, improving the interpretability of the fitted PRS models (**Supplementary Figure 6, Supplementary Table 3**). GWAS summary statistics-based methods tended to use upwards of 1 million variants either due to Bayesian modeling or large-scale ensembling. iPGS+refit, which enforces sparsity with Lasso penalized regression, consistently over-selected the number of variants, 5 times more than the true number of causal variants in the sparsest setting. On the other hand, SPLENDID only selected about 1.5 times the true number of causal effects for lower polygenicity and almost exactly the true support size for higher polygenicity. The much sparser model provided by SPENDID improves model interpretability.

Furthermore, SPLENDID use more GxPC interactions when training data was more diverse and heterogeneous (**Supplementary Figure 7, Supplementary Table 3**), showing that inclusion of more non-European ancestry is critical to accurately capturing effect heterogeneity. Even within a fixed sample size of 100,000, including more individuals of diverse ancestry into model training enabled better detection of heterogeneity in genetic effects. Using the estimated main and interaction effects to compute individual-level effects of selected variants (Equation 2), the resultant effect estimates were able to mirror the true effect sizes in each ancestry (**Supplementary Figure 8**). For more polygenic settings, the magnitude of heterogeneity was likely too weak for interactions to be selected, but larger sample sizes may enable SPLENDID to detect these signals. In these settings, we found that SPLENDID selected fewer than the true causal number of variants, whereas iPGS+refit selected slightly more, which may explain its slightly higher prediction accuracy.

Our simulation results showed that SPLENDID could consistently yield robust and higher prediction accuracy compared to existing approaches across various levels of polygenicity and genetic correlation structures.

### Analysis of quantitative traits in All of Us Research Program and UK Biobank

We compare SPLENDID with the existing approaches using data from AoU^25^ and UKBB^26^, two large biobanks with genotype and phenotype data for individuals of diverse ancestry (**Supplementary Table 4-5**). We analyzed 9 traits, including height; four lipid traits – low-density lipoprotein cholesterol (LDL), high-density lipoprotein cholesterol (HDL), triglycerides (log-TG), total cholesterol (TC); and four blood traits – neutrophil count (NEUT), white blood cell count (WBC), mean corpuscular hemoglobin (MCH), and MCH concentration (MCHC). Our analysis was restricted to unrelated individuals, and about 1.1 million variants in the AoU genotyping array data (version 7). We performed PCA using 1000G so that AoU and UKBB genotypes could all be projected onto the same PC space (**Methods**). We trained each method in AoU and conducted PRS tuning and validation in UKBB, using ancestry-specific GWAS summary statistics for PRS-CSx, CT-SLEB, and PROSPER, and individual-level data for iPGS+refit and SPLENDID (**Supplementary Table 6**). The UKBB data were randomly split in half for tuning and validation, where genetic ancestry was determined using random forest classification based on 1000G (**Supplementary Figure 9**).

Averaged across nine traits SPLENDID achieved similar or higher accuracy in non-European ancestry compared to all four alternative methods: a 82.3% higher adjusted R^2^ than iPGS+refit, 65.8% higher than PRS-CSx, 42.1% higher than CT-SLEB, and 27.8% higher than PROSPER (**Figure 3, Figure 4, Supplementary Tables 7-8**). In AFR specifically, SPLENDID had the highest adjusted R^2^ for all traits except for NEUT. Conversely, in EUR, SPLENDID performed worse than CT-SLEB and PROSPER in seven traits (from 2.7% in MCH to 23.2% in NEUT). This disparity is likely due to the much larger EUR sample size in the tuning data. For a more fair comparison and reducing health disparity in genetic research, when decreasing the EUR sample size to be similar with other ancestry groups when tuning in UKBB, these methods no longer outperformed SPLENDID in EUR for any trait (**Supplementary Figure 10**). This suggests that differences in PRS accuracy across ancestries can be influenced by the size of tuning data, particularly for methods like CT-SLEB and PROSPER that benefit from larger tuning sample sizes for optimal performance in specific ancestries. This also highlights the value of SPLENDID’s use of pooled ancestries in both training and tuning to provide more robust predictions across diverse populations without requiring any discrete labels.

**Figure 3.**
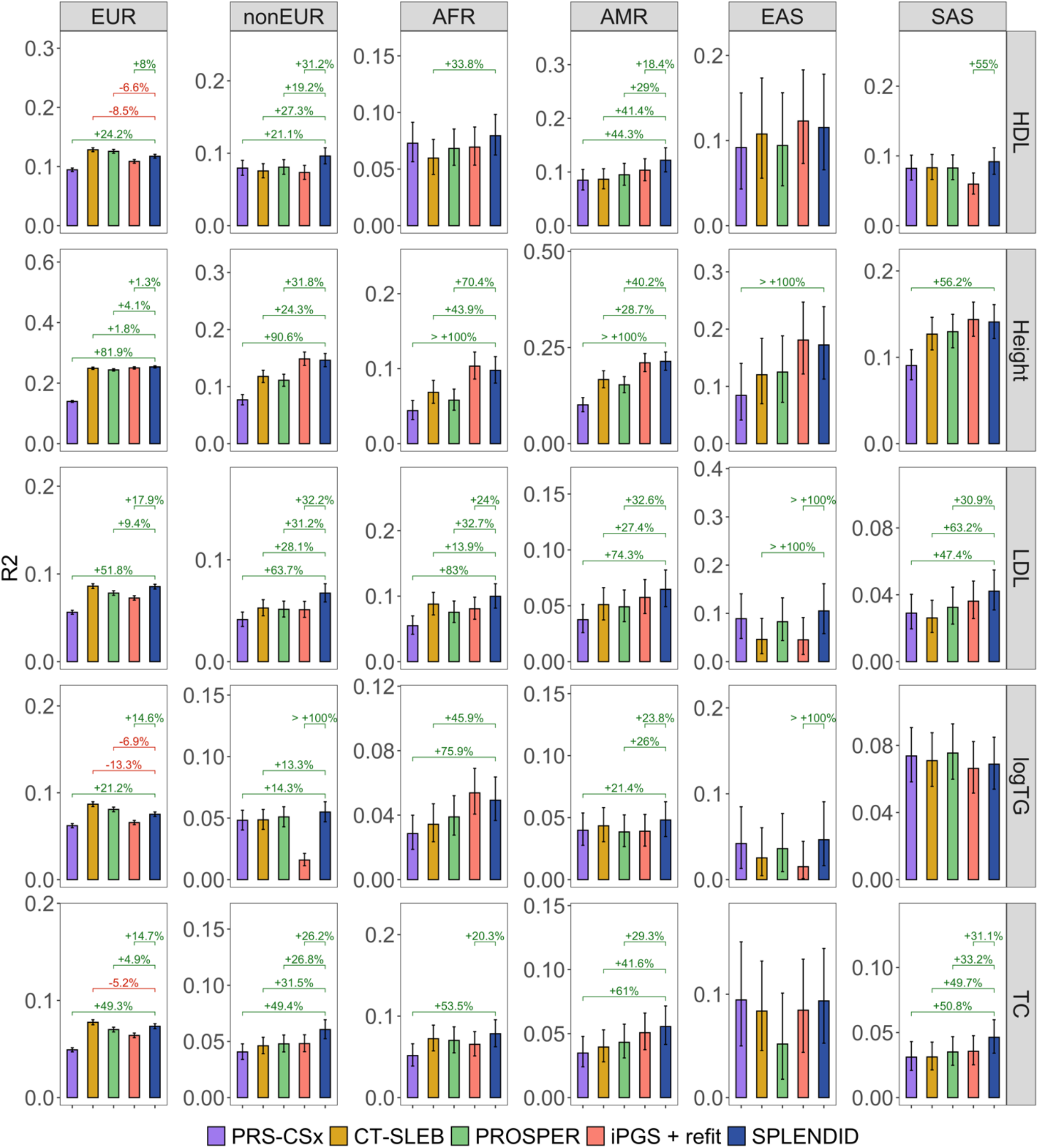
Prediction accuracy of PRS methods in four lipid traits and height. We trained each PRS method in AoU (n≈222,000), and used UKBB for tuning (n≈170,000) and validation (n≈170,000). Adjusted prediction R^2^ was reported as the correlation between the PRS and the residuals of each trait after regressing out age, sex, and the first 20 ancestry PCs. Error bars represent the 95% BCI of R^2^ from 10,000 bootstrap replicates. 95% BCI were also constructed for the percent increase in R^2^ by SPLENDID, where values are shown for results where this 95% BCI did not include 0.

**Figure 4.**
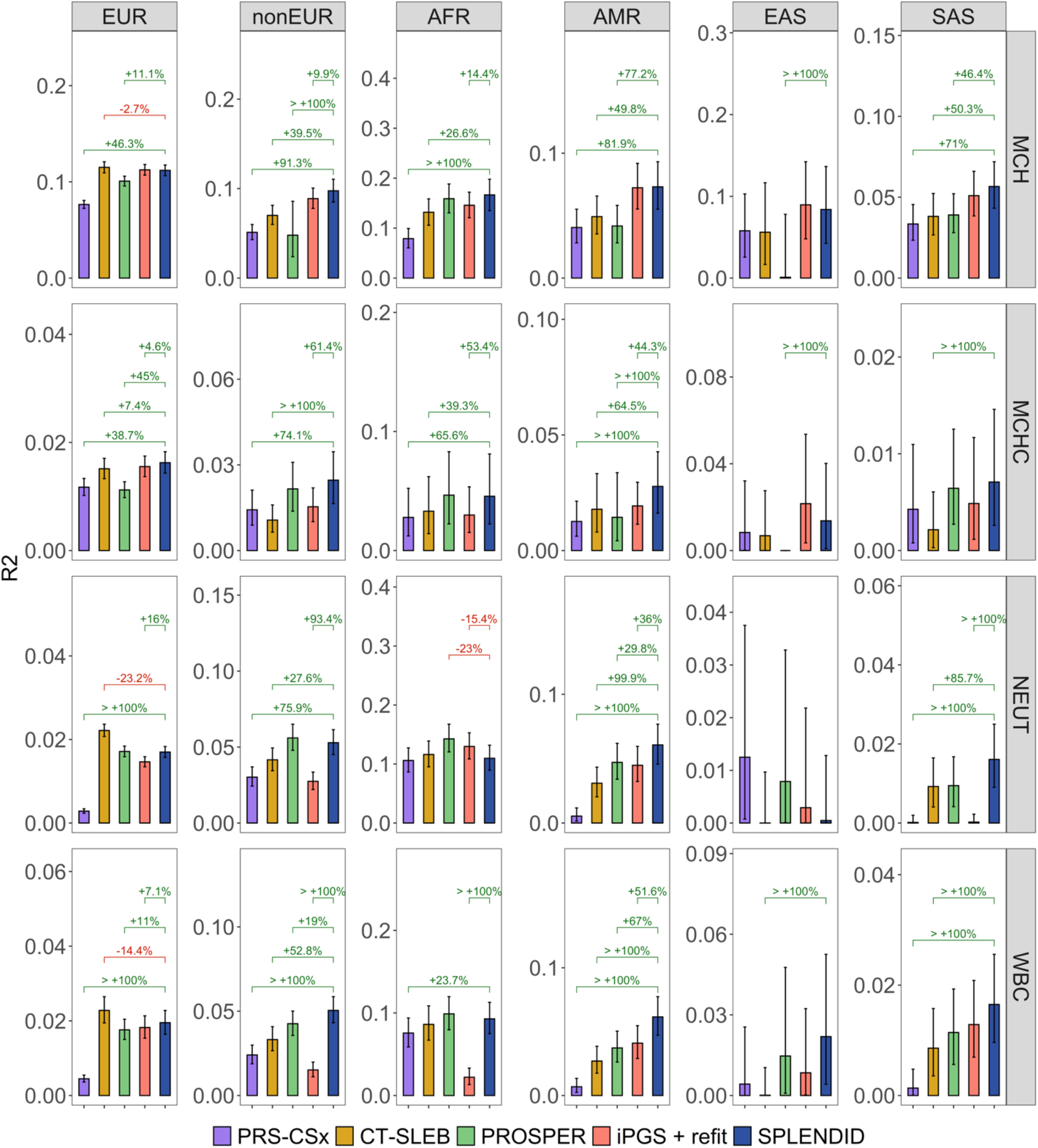
Prediction accuracy of PRS methods in four blood traits. We trained each of PRS method using AoU (n≈130,000) and used the same UKBB data for tuning and validation. Adjusted prediction R^2^ was reported as the correlation between the PRS and the residuals of each trait after regressing out age, sex, and the first 20 ancestry PCs. Error bars represent the 95% BCI of R^2^ from 10,000 bootstrap replicates. 95% BCI were also constructed for the percent increase in R^2^ by SPLENDID, where values are shown for results where this 95% BCI did not include 0.

In addition to improved prediction accuracy, SPLENDID consistently selected a substantially smaller number of predictive genetic variants compared to other methods (**Supplementary Table 9**). SPLENDID selected an average of 1,803 variants across the nine traits, less than half the average number of variants selected by iPGS+refit (3,959). GWAS summary statistics-based methods used significantly more – between 300,000 and 450,000 variants for PRS-CSx and CT-SLEB and around 950,000 for PROSPER.

Overall, SPLENDID yielded fairly consistent improvements in prediction accuracy over existing methods, particularly in non-European ancestries. Furthermore, by only using a concise set of predictive variants, our method can lead to easier interpretation of individualized genetic risk, better identification causal variants, and fairer implementation in clinical settings.

### Trait-specific allele frequency and effect size differences

The three key challenges of PRS transportability between ancestries are LD, allele frequency, and effect size differences. The first two can be addressed by pooled analysis across diverse ancestries. Inclusion of individuals across diverse backgrounds leverages power from populations with higher allele frequencies for predictive variants^27^, and may identify variants that are more likely to be causal rather than ancestry-specific tagging variants from different LD patterns^9,28–30^. Specifically, SPLENDID selects several variants that are common (MAF > 1%) only in non-European ancestries (**Supplementary Figure 11, Supplementary Table 10**). This highlights the value of pooled analysis across multiple ancestries in identifying predictive variants that may not be detectable from analyzing a single ancestry, such as EUR. One clear example is the Duffy-null variant (rs2814778) on chromosome 1 that is attributed to lower white blood cell counts in healthy individuals of African ancestry^31–33^. This variant has MAF above 10% in AFR and AMR within AoU, but it likely would not be selected if the analysis was restricted to EUR only.

The main innovation of SPLENDID is the use of GxPC interactions to model effect heterogeneity across ancestries by treating ancestry as a continuum rather than distinct categories. We found that traits such as LDL and HDL showed very similar effect sizes across ancestries, which may suggest that AF differences are a key driver of varying PRS accuracy (**Supplementary Figure 11** and **Supplementary Figure 12**). However, we observed greater heterogeneity in effect sizes for blood cell traits, particularly between EUR and AFR (**Supplementary Figure 12**). For example, in our analysis of MCH, SPLENDID selected a variant on chromosome 16 (rs1211375) with allele frequencies well above 20% in all ancestries but showed particularly strong individual-level effect sizes in AFR (**Figure 5**), which aligns with existing literature^34,35^, showing that our method indeed identifies biologically relevant variants with heterogeneity.

**Figure 5.**
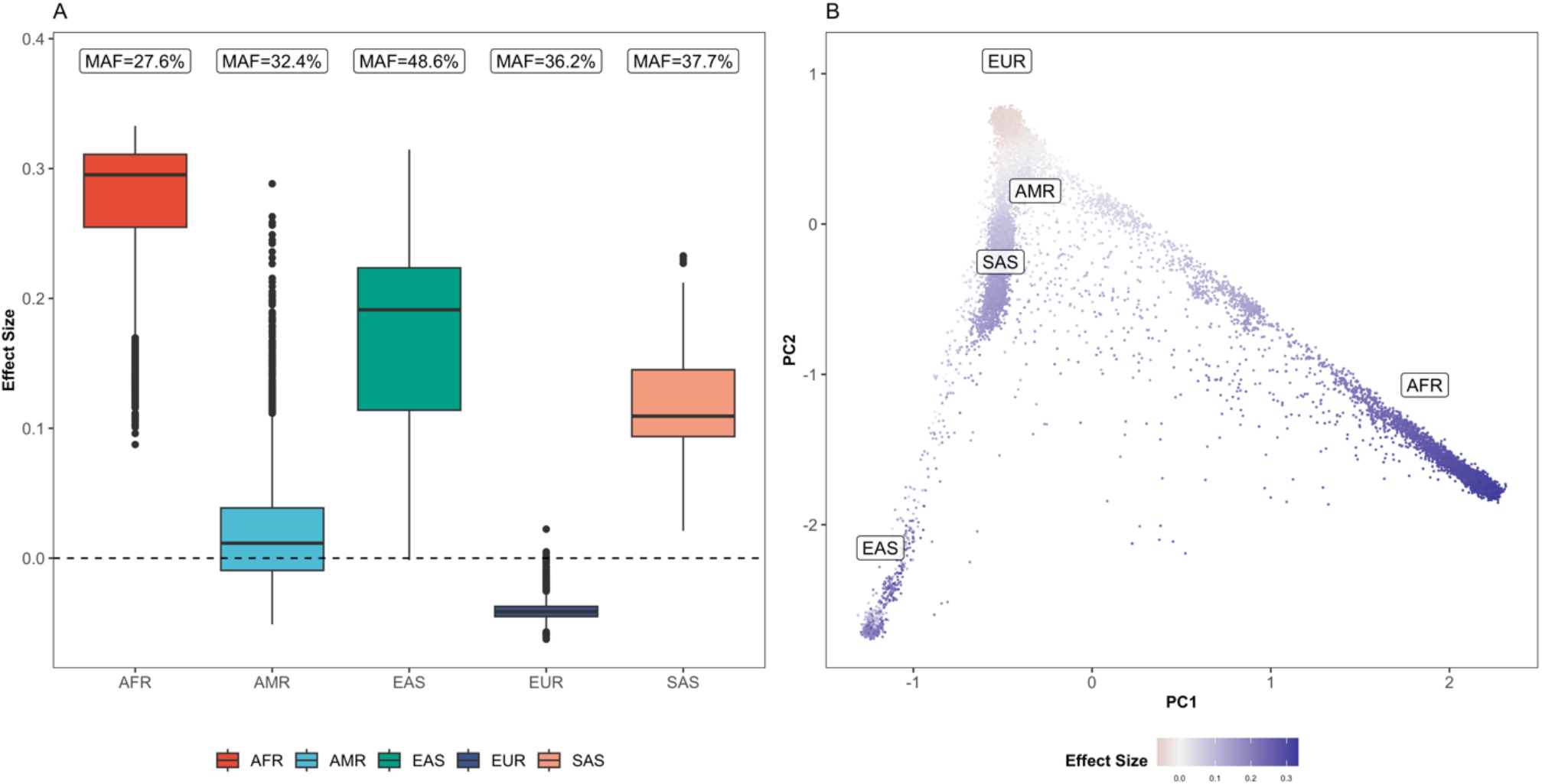
Individual-level effects of rs1213375 on mean corpuscular hemoglobin across genetic ancestry. The rs1213375 variant was selected by SPLENDID with strong interaction effects for MCH. We computed individual-level effect sizes using estimated main and interaction effects and the first 5 ancestry PCs in the UKBB validation dataset. (A) We compare the distribution of individual-level effect sizes in different ancestry groups, with text boxes showing the MAF of the variant for each ancestry. (B) We also show this distribution among all samples across the continuum of genetic ancestry, quantified by the first two ancestry PCs (standardized to mean 0 and variance 1 for visualization). Text boxes indicate regions of the PC space roughly corresponding to each genetic ancestry group.

By leveraging pooled-ancestry analysis and incorporating GxPC interactions into the model in constructing PRS using diverse biobank-scale data, SPLENDID identified several variants that may address challenges of PRS transportability due to ancestry differences in MAF, LD, and effect size. While most selected interactions do not have published evidence for heterogeneity like rs1211375, a sparse set of selected variants and GxPC interactions provides an initial search space to further investigate biological mechanisms underlying effect heterogeneity.

### Testing for heterogeneity with meta-analysis of published GWAS

In our primary model design, we used GxPC interaction multi-ancestry GWAS in the training data to screen for potential effect heterogeneity. However, depending on data availability and computing resources, it may be more convenient or advantageous to use published GWAS summary statistics, which often have much larger sample sizes than a single biobank. To evaluate this, we used METAL^36^ to conduct meta-analyses of summary statistics from the Global Lipid Genetics Consortium (GLGC)^10^, Genetic Investigation of Anthropometric Traits (GIANT) consortium^37^, and Blood Cell Consortium (BCX)^38^. We tested for effect size heterogeneity between ancestries using the Q-statistics from each meta-analysis.

Across nearly all traits and ancestries, there was no consistent difference in adjusted R^2^ between the two screening approaches (**Supplementary Figure 13, Supplementary Table 11**). Using external GWAS meta-analysis yielded significant improvements in adjusted R^2^ for HDL and TG in AMR (4.3% and 11.7%, respectively), as well as NEUT and WBC in AFR (7.6% and 11.9%, respectively), but a significant 5.6% decrease in adjusted R^2^ for HDL in SAS and a mix of results in EUR (from -7.6% to +5.9%). While results varied between traits and ancestries, these results suggest that incorporating external GWAS data within the SPLENDID framework may improve prediction accuracy by more accurately identifying relevant heterogeneous genetic effects.

### Robustness to ancestry proportions in UKBB tuning data

We conducted additional sensitivity analysis to evaluate SPLENDID’s prediction accuracy given different proportions of EUR ancestry in the tuning dataset by down-sampling EUR samples. The original UK Biobank tuning data has over 150,000 EUR individuals out of about 160,000 total independent samples (over 90%). We subsampled the tuning data to have only 5,000 of EUR (about 30%), which is more comparable to the sample sizes for AFR, AMR, and SAS. Across the nine traits, we found that SPLENDID attained similar adjusted R^2^ with either tuning dataset. The more balanced ancestry proportions in the down-sampled tuning data yielded slight improvements, primarily in African ancestry (**Supplementary Figure 14, Supplementary Table 12**). Simultaneously, there were significant decreases in adjusted R^2^ in European ancestry. However, these trends were not uniform; for instance, tuning with down-sampled EUR yielded a 1.1% decrease in adjusted R^2^ in EUR and 9.5% increase in AFR for HDL, a 0.9% increase in EUR and 17.4% increase in AFR for logTG, and a 6.2% decrease in EUR and 6.9% decrease in AFR for TC. In over half of the trait-ancestry combinations, there was no significant difference in R^2^ using either ancestry proportion.

We found that SPLENDID was robust to the ancestry proportions of the UKBB tuning data, whether it was more balanced or skewed towards EUR. This stands in contrast to methods like CT-SLEB and PROSPER, whose accuracies dropped below that of SPLENDID after decreasing the EUR tuning sample size (**Supplementary Figure 10**). This is a critical step in moving towards label-free analysis to improve prediction accuracy without major concern about the particular ancestry makeup of the data.

### Cross-fold training for prediction modeling without tuning data

Our primary framework requires of an independent tuning dataset for ensemble learning. However, we can also build a prediction model with only training data. Following the previously proposed cross-model selection and averaging (CMSA) approach from bigstatsr^39^, we ran five-fold group-L0L1 regressions in the AoU training data, averaging together the best models from each fold to produce a final prediction model. Using the same UKBB validation dataset as in our main analyses, we compared this SPLENDID-CMSA approach with PRS-CSx-meta-auto, a fully Bayesian version of PRS-CSx that automatically selects prior hyperparameters and combines posterior effect estimates from each ancestry.

Using LDL and MCH as illustrative examples, we found that SPLENDID-CMSA significantly outperformed PRS-CSx-meta-auto overall and each ancestry except SAS for MCHC (**Supplementary Figure 15, Supplementary Table 13**). SPLENDID-CMSA outperformed PRS-CSx-meta-auto by over 100% for LDL and 74.7% for MCH, while also achieving 38.3% and 31.8% higher adjusted R^2^ than the original PRS-CSx for LDL and MCH, respectively. However, SPLENDID-CMSA had about 10% lower adjusted R^2^ compared to the main implementation with ensembling in UKBB. While ensemble learning in external tuning data generally performed better in our analyses, we have demonstrated that SPLENDID can achieve comparable accuracy when tuning data is not readily available, and other cross-model approaches could be explored to further enhance accuracy in this scenario.

## Discussion

In this study, we propose SPLENDID, a novel method for PRS prediction that leverages biobank-scale genetic and clinical data across diverse populations. By treating ancestry as a continuum and incorporating individuals from all ancestries in both the model training and tuning processes, SPLENDID constructs a single, generalizable prediction model. This approach eliminates the need to develop separate PRS models for different ancestry groups, enabling more inclusive and accurate predictions and increasing the fairness in clinical settings.

Through extensive simulation and real-world applications to multi-ancestry biobanks, such as AoU and UKBB, SPLENDID demonstrated robust and higher prediction accuracy compared to state-of-the-art PRS methods. By training on diverse data and incorporating ancestry-dependent genetic effects through GxPC interactions, SPLENDID captures important differences in genetic factors like MAF, LD, and effect size across ancestries populations. In traits such as neutrophil count and MCH, SPLENDID identified several biologically relevant variants, highlighting its potential to offer deeper insights into the genetic architecture of complex traits and diseases.

One of SPLENDID’s strengths is its robustness to different ancestry compositions within tuning datasets and its ability to maintain high accuracy even without tuning data by using a cross-model strategy that relies solely on training data. This feature is especially beneficial for predicting common diseases, where large-scale data from disease consortia can be used to improve prediction accuracy^40–45^.

However, there are some limitations to this study that open opportunities for future development. Our analysis focused on genetic interactions with ancestry PCs based on 1000G given its wide availability, where new validation genotypes can be easily projected to the same PC space. However, 1000G data may not capture the full spectrum of human genetic diversity, particularly individuals with admixed ancestry. Developing more globally diverse reference data^46^ and incorporating other notions of genetic ancestry besides PCs^30,47–49^ to capture and understand differences in genetic risk across ancestries. Moreover, while genetic ancestry is a crucial factor, it alone may not fully address the challenges of PRS accuracy in diverse populations. To that end, the SPLENDID framework and future extensions can also be used to model complex genetic interactions with clinical, social, and environmental factors to more comprehensively model genetic risk^50–53^.

In addition, the SPLENDID framework is limited to analysis of individual level data from multi-ancestry biobanks, like AoU. In contrast, methods such PRS-CSx^12^ and CT-SLEB^15^ can use summary statistics from large-scale GWAS meta-analyses. While SPLENDID outperformed these GWAS-based methods using the same training data, many published GWAS have substantially larger sample sizes and potential for greater predictive power. Thus, our framework can be expanded in the future to integrate external data such as published GWAS and global biobanks^54^, which will involve more advanced model designs and larger efforts in cloud computing and data harmonization^54–57^.

In conclusion, SPLENDID offers a new approach to polygenic risk prediction by modeling ancestry on a continuous scale and leveraging large-scale, diverse datasets. Our analysis shows that SPLENDID can uncover the genetic factors underlying complex traits, providing a more precise and inclusive framework for genetic risk prediction. With the expansion of genetic and clinical data from both biobanks and consortia, SPLENDID stands as a promising tool for more accurate and personalized risk prediction across diverse populations, increasing its clinical practical utility.

## Acknowledgements

We would like to thank Rui Duan and Peter Kraft for their discussions, feedback, and insights on this work. We would also like to thank Xihao Li, Derek Shyr, Hufeng Zhou, Kangcheng Hou, Florian Privé, Xing Hua, Jacob Williams, and Wendy Wong for guidance and assistance with software development and the All of Us platform. Analysis for this work utilized the high-performance computation Biowulf cluster at the National Institutes of Health (NIH) and Faculty of Arts and Sciences Research Computing Cluster at Harvard University. Finally, we are grateful for the participants in the 1000 Genomes Project, UK Biobank (obtained from resource application no. 52008) and All of Us Research Program for providing vital data to this project.

This research was supported by NIH Training Grant T32GM135117 and NSF Graduate Research Fellowship DGE-2140743 (T.C.), NIH Intramural Research Program (H.Z.), Office of Naval Research grants N000142112841 and N000142212665 (R.M.), and NIH grants R35-3 CA197449, U19-CA203654, R01-HL163560, U01-HG009088, U01-HG012064 (X.L.).

The All of Us Research Program is supported by the NIH, Office of the Director: Regional Medical Centers: 1 OT2 OD026549; 1 OT2 OD026554; 1 OT2 OD026557; 1 OT2 OD026556; 1 OT2 OD026550; 1 OT2 OD 026552; 1 OT2 OD026553; 1 OT2 OD026548; 1 OT2 OD026551; 1 OT2 OD026555; IAA no.: AOD 16037; Federally Qualified Health Centers: HHSN 263201600085U; Data and Research Center: 5 U2C OD023196; Biobank: 1 U24 OD023121; The Participant Center: U24 OD023176; Participant Technology Systems Center: 1 U24 OD023163; Communications and Engagement: 3 OT2 OD023205; 3 OT2 OD023206; and Community Partners: 1 OT2 OD025277; 3 OT2 OD025315; 1 OT2 OD025337; 1 OT2 OD025276.

## Data and Software Availability

Software and tutorials for SPLENDID are available on GitHub (https://github.com/chen-tony/SPLENDID). Simulated multi-ancestry genotype data is available on the Harvard Dataverse (https://dataverse.harvard.edu/dataverse/multiancestry). Data from 1000G were obtained from https://www.cog-genomics.org/plink/2.0/resources (genotypes) and https://www.internationalgenome.org/data-portal/sample (population labels)

## Methods Model setup

We first assume a linear model of an outcome *Y* ∈ ℝ^*n*^ as a function of genotypes *G* ∈ ℝ^*n*×*M*^ with *M* total genetic variants, *K* ancestry principal components *PC* ∈ ℝ^*n*×*K*^, and *L* baseline covariates *Z* ∈ ℝ^*n*×*L*^. For an individual *i* ∈ {1 … *n*}, we assume genetic variant *j* ∈ {1 … *M*} has effect size *B*_*ij*_, PC *k* has effect *γ*_*k*_, and covariate *m* has effect *α*_*m*_, with an error term *ϵ*_*i*_:

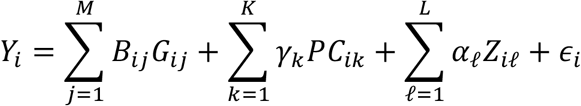

The genetic effect sizes *B*_*ij*_ are assumed to be either homogeneous for all individuals or heterogeneous across ancestries, which can be characterized by the first *K*_0_ ≤ *K* ancestry PCs:

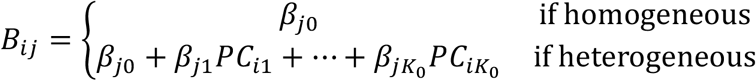

Denoting *H* as the set of variants *j* with heterogeneous effects, we can expand the first linear model into a GxPC interaction model.

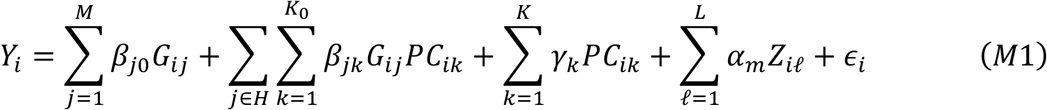

For our analyses, we used the first *K* = 20 ancestry PCs for baseline covariates, and the first *K*_0_ = 5 ancestry PCs for interactions. Using these five PCs can be roughly interpreted as accounting for the five broadly-defined continental populations, thereby capturing the major axes of genetic variation.

### Principal components analysis

We used 1000G as a universal reference for PCA to ensure consistency across training, tuning, and validation datasets. We first performed LD pruning of all variants using the PLINK 1.9^58^ command `--indep-pairwise 500 5 0.05`, which creates a reduced subset of relatively independent variants. This step is crucial to mitigate the effects of correlated variants to improve the robustness of PCA.

After LD pruning, we computed the first 20 PCs using the PLINK 2.0^59^ command `–pca 20 allele-wts`, which provides the corresponding PC loadings. These loadings are essential as they allow for project of new data onto the same PC space defined by the reference dataset.

To standardize PCs, we rescaled them to have a mean of 0 and variance of 1 based on their distribution in the training data. This standardization ensures that PCs are on a comparable scale and can be effectively used as covariates and interaction terms in subsequent analyses. It is important to note that pre-computed in-sample PCs, such as those provided by AoU and UKBB, do not align with our reference PCs. These study-specific often lack the necessary PC loadings to project new data to the same PC space, which could lead to discrepancies when applied to external datasets.

### Screening for heterogeneity

To reduce the number of interactions in the model, we run a GWAS analysis with GxPC interactions in the training data. For each variant *j*, we run a marginal regression of the following form:

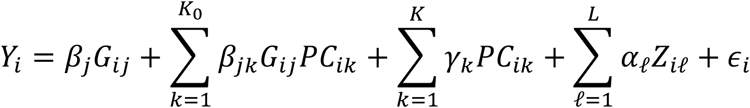

Then, we compute F-statistics to test the null hypothesis that none of the *K*_0_ GxPC interactions are associated with the phenotype:

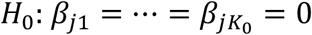

We conduct this analysis using PLINK 2.0^59^, using the highlighted commands for our particular model:

~~~
∼/software/plink2 --bfile BFILE --linear **interaction** hide-covar --pheno
PHENO.txt --pheno-name --covar COV.txt --covar-name
age,sex,pc1,pc2,pc3,pc4,pc5,pc6,pc7,pc8,pc9,pc10,pc11,pc12,pc13,pc14,pc15,pc1
6,pc17,pc18,pc19,pc20 **--parameters 1-23**,**26-30 --tests 24-28** --out OUT
~~~

The inputs for the “--parameters” flag indicate the main effects of the variant and all covariates (1-23), and the interactions between the variant and the first 5 PCs (26-30), skipping interaction terms with age (24) and sex (25). The inputs for the “--tests” flag call for a joint test for the 5 GxPC interactions (24-28 after skipping interactions with age and sex).

We use the corresponding F-test p-values *p*_*j*_ of each variant and compared against various p-value thresholds 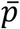 to identify those with potential heterogeneous effects across ancestries. For those variants, we included GxPC interactions in the subsequent penalized regression.

### Group-L0L1 penalized regression

To select a concise set of predictive genetic variants and potential GxPC interactions, we use a group-L0L1 penalized regression framework. The group L0 penalty corresponds to the number of groups selected, where each group consists of a genetic variant and any GxPC interaction terms. The group L1 penalty refers to the sum of L2-norms for each group, which shrinks coefficients of selected groups for improved out-of-sample prediction and can be thought of as an extension of the L1 penalty from traditional Lasso regression to the group selection context (i.e. group Lasso)^60^. We chose this particular penalized approach rather than the group Lasso because the L0 penalty component enables even sparser group selection without sacrificing prediction accuracy^22^. This allows for better interpretation of the prediction model by focusing on a concise set of variants to understand the key drivers of an individual’s genetic risk, as well as more convenient implementation in practice by reducing the number of variants to process and analyze.

Group-L0L1 penalized regression aims to optimize the following objective function

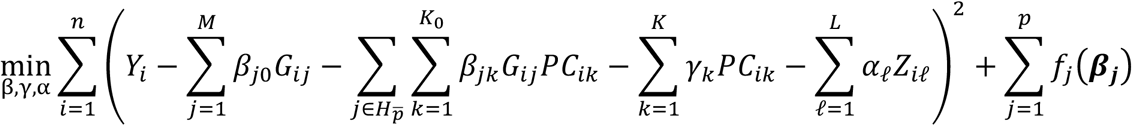

where ***Y*** and the columns of ***PC*** and ***Z*** are all standardized to have mean 0 and variance

1. Using a given p-value threshold 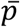, for variants with GxPC p-value 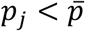.we apply a grouped L0L1 penalty to jointly penalize the main and interaction terms 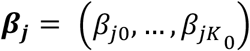.For variants with 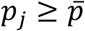,we only penalize the main effects *β*_*j*0_, which reduces to a simple L0L1 penalty. Thus, the penalty function *f*_*j*_(***β***_***j***_) is defined as

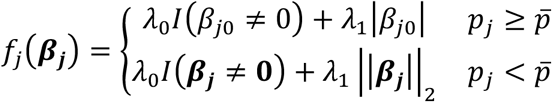

where 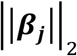 is the L2 norm defined as 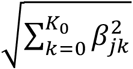 for the vector of main and interaction effects, the sum of which corresponds to the group L1 penalty.

This optimization problem can be computationally challenging, particularly with large-scale and high-dimensional data, but one can obtain high-quality solutions at scale by using a cyclical coordinate descent-based approach along with active set updates^22^. At each iteration *t* and variant *j*, we update residuals *r*_*i*_ as the outcome minus the overall fitted value based on the current estimates for ***β, γ***, and ***α***. For variants *j* ∈ *H* define a matrix 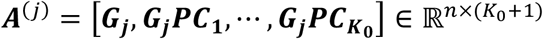, which concatenates the genotypes for variant *j* and the *K*_0_ GxPC interactions. For variants *j* ∉ *H*, we simply let ***A***^(*j*)^ = [***G***_***j***_ ] ∈ ℝ^*n*×1^. For each variant, we then define *L*_*j*_ as the largest eigenvalue of 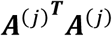. Then, the coordinate update for ***β***_***j***_ is given by

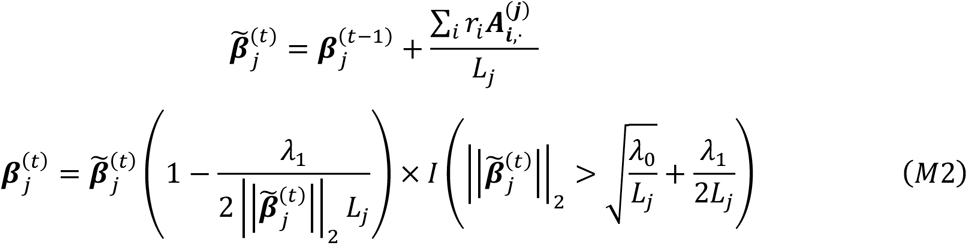

This regression framework requires individual-level genotype and phenotype data. We use software from the bigstatsr and bigsnpr packages^61^ for memory-efficient handling of large genotype datasets by converting PLINK-format files into file-backed matrix format^39,61,62^.

Within our optimization algorithm, we employ active set updates to minimize the total number of processed during each iteration of coordinate descent. At each iteration *t*, we only loop through the indices *j* with nonzero *β*_*j*0_ in the previous iteration *t* − 1. This approach significantly reduces runtime, particularly when genetic effects are sparse, by excluding variants don’t contribute much to the trait. Once the active set converges, we cycle through the excluded variants to identify any that might further reduce the loss. These variants are reintegrated into the active, and coordinate descent is rerun to ensure stability of the final solution.

For parameter tuning, we use three values of 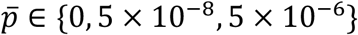,where 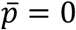includes no interactions in case a given trait has minimal effect heterogeneity. For penalization, we start with five values of *λ*_1_ on the log-scale from 10^−2^ to 10^−4^. For each value of *λ*_1_ we compute an initial L0 penalty parameter

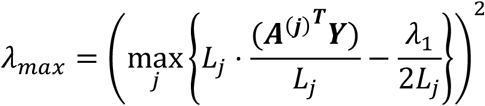

which corresponds to the smallest *λ*_0_ where no variants would be selected. Using this *λ*_*max*_, we consider up to 50 values of *λ*_0_ starting from 0.95*λ*_*max*_ and decreasing down to 0.0005 *λ*_*max*_ on the log scale. Borrowing from the bigstatsr implementation of Lasso regression^39^, for each *λ*_1_ value, we allow the algorithm to stop along the *λ*_0_ path when the squared error loss in an external tuning dataset stops decreasing, or when the total number of selected variants reaches a certain upper bound. These features avoid excess computation of candidate models that either do not offer much prediction power or include too many variants and can be adjusted based on preferred features of the prediction model and computational resources.

### Sparse ensemble learning

Ensemble learning is a powerful technique widely used to enhance the prediction accuracy of PRS^5,14–16,23,63^. However, traditional ensembling methods often incorporate a large number of variants in the final PRS, which will hinder interpretability and practical implementation. To address this, we employ a sparsity-aware approach to maintain sparsity in the final PRS while leveraging the benefits of improved accuracy from ensembling.

In our procedure, we first compute the R^2^ of PRS from each tuning parameter combination 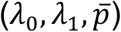 within the tuning dataset to identify a single best PRS, as well as the corresponding set of selected variants *S*. We then construct a set of “ensembling candidates” by iterating through all PRS estimates ordered by tuning R^2^ and incrementally adding PRS such that the total number of variants included is less than *c*|*S*|, where *c* is a desired multiplicative factor. Our default choice is *c* = 2, allowing us to use at most twice the number of variants selected by the best single PRS, but this can be modified depending on sparsity interests. This factor *c* can be adjusted based on specific sparsity targets.

Next, we use cross-validated ridge regression of the phenotype in the tuning data against the chosen ensembling candidates to produce an optimal weighted combination of all candidate PRS. This approach ensures that we achieve a balance between sparsity and prediction accuracy, enhancing the interpretability and practical utility of the final PRS.

### Simulation analysis

We designed our simulation analyses based on simulated genotype data previously generated and described in the CT-SLEB paper^15^. Briefly, the data included a total of 600,000 independent samples with reference LD based on 1000G. There were 120,000 samples representing each of five continental ancestries (AFR, AMR, EAS, EUR, and SAS). We generated effect sizes from a mixture distribution with the probability *π* of having nonzero effects, where the non-zero effect sizes follow five-dimensional multivariate normal distributions:

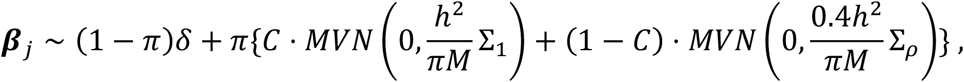

where *M* is the total number of variants, *δ* is a point mass at 0, *π* is the proportion of variants with nonzero true effect size, Σ_*ρ*_ is a five-by-five correlation matrix (one column and row for each population) with off-diagonal entries *ρ*, Σ_1_ is a matrix of 1’s, and *C* is a binary variable that denotes whether the variant has identical effect sizes across all five populations or varies with correlation *ρ*. For our simulations, we use *π* =0.05%, 0.5% and 1% with *h*^2^ =0.4 and *M* =1,158,251. We then used PLINK 2.0 to simulate 10 replicates for each setting. To simulate phenotypes, we multiplied genotypes by causal effects for a baseline genetic score (*Gβ*). Then, for each simulation replicate, we added normally-distributed random noise to achieve a total heritability of approximately 0.4.

For all methods, we used a fixed training sample size of 100,000 with varying ancestry proportions. We considered non-European proportions of 0%, 30%, 50%, and 80%, with equal proportions among the four non-European ancestries (i.e. AFR, EAS, AMR, and SAS). We used 18,000 samples (10,000 European and 2,000 of each non-European ancestry) for tuning and 50,000 samples for validation (10,000 for each ancestry).

iPGS+refit and SPLENDID were trained and tuned using all ancestries combined, where we stopped the parameter grid search after the squared error in the tuning data stops decreasing. iPGS+refit also has an ancestry-specific refit step. PRS-CSx, CT-SLEB and PROSPER were trained used ancestry-specific GWAS summary statistics from the training samples and were tuned separately in each ancestry.

### Real data analysis in All of Us Research Program and UK Biobank

We used phenotype, covariate, and genotype array data from AoU (version 7) to train each method, capitalizing on diverse patient population. Tuning and validation were conducted in the UKBB, which is less ancestrally diverse but still contains a substantial non-European sample size for method comparison across different ancestries.

We analyzed nine continuous traits: LDL, HDL, log-TG, TC, height, neutrophil count, WBC, MCH, and MCHC. In AoU, these traits were extracted from electronic health records, using the most recent measurements for each individual. We restricted our training data to unrelated individuals with short-read whole-genome sequencing using available quality-control information from AoU Research Platforms. Detailed information about genotyping, quality control measures, and the methodologies for excluding related participants were thoroughly documented in the All of Us Research Program Genomic Research Data Quality Report. In UKBB, we used UKBB Field Codes to extract measurements, and subsetted to unrelated individuals up to 3^rd^ degree relatives. For individuals with reported statin usage, we divided LDL cholesterol measurements by 0.7 and total cholesterol by 0.8, per standard practices^64^.

For variant selection, we started with the All of Us Global Diversity Array, which has around 2 million variants, and subsetted to the intersection with UK Biobank and 1000 Genomes Project Phase 3 genotypes. We then filtered for variants with minor allele frequency above 1% for at least one of the six predicted ancestry groups in AoU (AFR, AMR, EAS, EUR, SAS, and MID for Middle East ancestry), giving a final set of 1,172,070 variants.

For the UKBB data, we classified individuals into one of the five major continental ancestries (AFR, AMR, EAS, EUR, SAS) for tuning of GWAS-based methods as well as ancestry-stratified validation results. First, within the 1000G reference dataset, we trained a random forest classification model using the first 20 ancestry PCs and corresponding continental population labels. We then applied this classification model to the first 20 PCs within the UKBB (projected to the 1000G PC space) to get predicted ancestry labels (**Supplementary Figure 9**). It is important to note that there were individuals with low predicted probabilities that were still assigned to one of the five ancestry categories. However, these labels were not required for any key step of SPLENDID, and were only necessary for ancestry-specific tuning of the existing PRS methods, as well as stratified validation results to compare between each method.

PRS-CSx, CT-SLEB, and PROSPER were conducted by computing GWAS summary statistics within each genetically-predicted ancestry group, and using 1000G for the LD references. We combined individuals of MID (Middle East) and EUR ancestries, since these methods do not have LD references for MID. For iPGS+refit and SPLENDID, we stopped the parameter grid search once the number of selected variants exceeded 10,000 to limit computation on the AoU platform. Training of all methods in AoU controlled for age, sex, and the first 20 PCs based on 1000G. Adjusted R^2^ of PRS in UKBB were computed with respect to residuals of each outcome after regressing out age, sex, and the first 20 PCs based on 1000G.

### Implementation of existing methods

*PRS-CSx*^*12*^ is a Bayesian approach that uses a continuous shrinkage prior to jointly model effect sizes using GWAS summary statistics and LD from multiple populations, where for variant *j* and population *k*, the effect size follows the prior

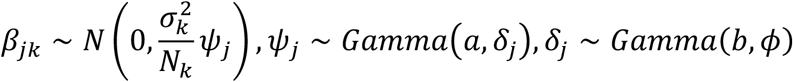

*ϕ* and *ψ*_*j*_ are global and local shrinkage parameters, respectively, to model genetic architecture, *σ*_*k*_ is the residual variance for population *k*, and *N*_*k*_ is the sample size of the GWAS for population *k*. The default values of *a* and *b* were 1 and ½, respectively. We performed grid search among *ϕ* values of 1, 10^−2^, 10^−4^, and 10^−6^ by finding the best linear combination of estimated effects within each of the five continental populations. For both simulation and real data analysis, we use pre-computed LD based on 1000G, as provided on GitHub.

*CT-SLEB*^*15*^ is a clumping-based method that implements 2-dimensional clumping and thresholding using summary statistics from a source population (i.e. European) and a target population (e.g. African). The method includes Empirical Bayes estimation of cross-ancestry effects, and ensembling using SuperLearner to optimize PRS in the target population. We used default parameter settings for LD threshold *r*^2^, window base *w*_*c*_, and p-value thresholds *p*_*T*_, with the exception of replacing p-value thresholds 0.5 and 1 with 0.3 to reduce computational demands. We conducted 2-way empirical Bayes estimation between European and each non-European ancestry, and included PRS estimates from all five ancestries in the SuperLearner step, using Lasso and Ridge learners. We used the 1000G, separated into each of the five continental ancestries, for LD clumping.

*PROSPER*^*16*^ is a penalized regression method that uses an elastic net penalty to model effects across ancestries. First, Lassosum2^65^ is run within each population *k* using the following penalized regression:

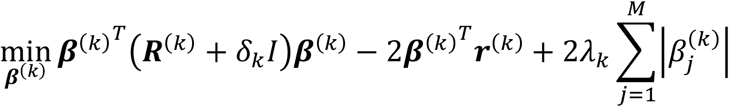

where ***r***^(*k*)^ and ***R***^(*k*)^ denotes the GWAS summary statistics and LD matrix for population *k*, respectively, and *λ*_*k*_ and *δ*_*k*_ are L1 and L2 penalty parameters, respectively. An optimal 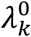 and 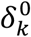 are selected for each population using grid search, and then input into the main penalized regression for cross-ancestry modeling:

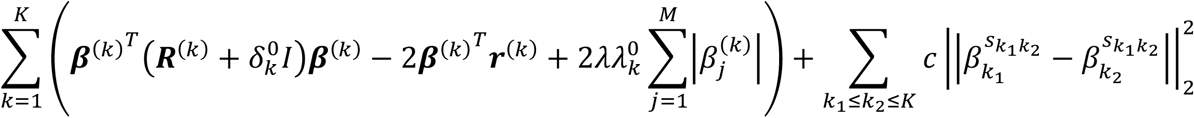

where *λ* models sparsity and *c* models shared effect sizes between pairs of populations, 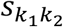 is the shared support (i.e. variants) between each pair of populations *k*_1_ and *k*_2_, and 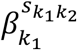 is the vector of effects for population *k*_1_ on that shared support. Using estimates from all tuning parameters and ancestries, we run SuperLearner with Lasso and Ridge learners within each ancestry. We omitted the OLS learner, which is included in the original software, as we found this component failed in certain traits and ancestries and resulted in nearly zero R^2^. For both simulation and real data analysis, we use pre-computed LD based on 1000G, as provided on GitHub.

*iPGS+refit*^*21*^ is a two-step penalized regression approach using individual-level data. The first step uses Lasso penalized regression to capture shared effects across ancestry

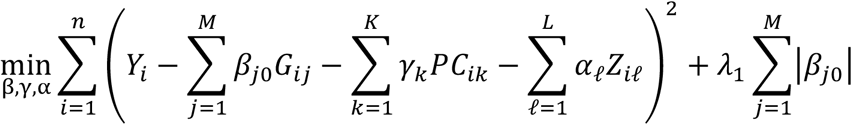

We implemented Lasso regression using the same optimization algorithm as SPLENDID, with *λ*_0_ = 0 and no GxPC interactions (i.e. *H* = ∅), and a log-scale grid of 100 *λ*_1_ values starting from 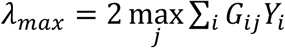 and stopping at *λ*_*min*_ = *λ*_*max*_ × 0.001.

Using the tuning dataset, we selected a fixed “iPGS” prediction model using the best *λ*_1_. In the second “refitting” step, we test for heterogeneity in variant effects based on meta-analysis of ancestry-specific GWAS from the training data using METAL^36^. Following the published implementation, we selected a set *H* as variants with heterogeneity p-values below 5 × 10^−8^. Then, we use the glmnet^66^ package to run cross-validated Lasso in each ancestry with the following regression model:

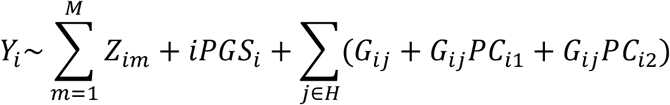

This step is designed to model ancestry-specific effects not captured by the main iPGS model. Following the original paper, we used a penalty factor (i.e., multiplicative factor on ***λ***_**1**_ for different variables) of 1 for iPGS, 1.1 for the main effects in ***H***, and 1.2 for interaction terms.

