## Supplementary Figures for "SPLENDID incorporates continuous genetic ancestry in biobank-scale data to improve polygenic risk prediction across diverse populations"

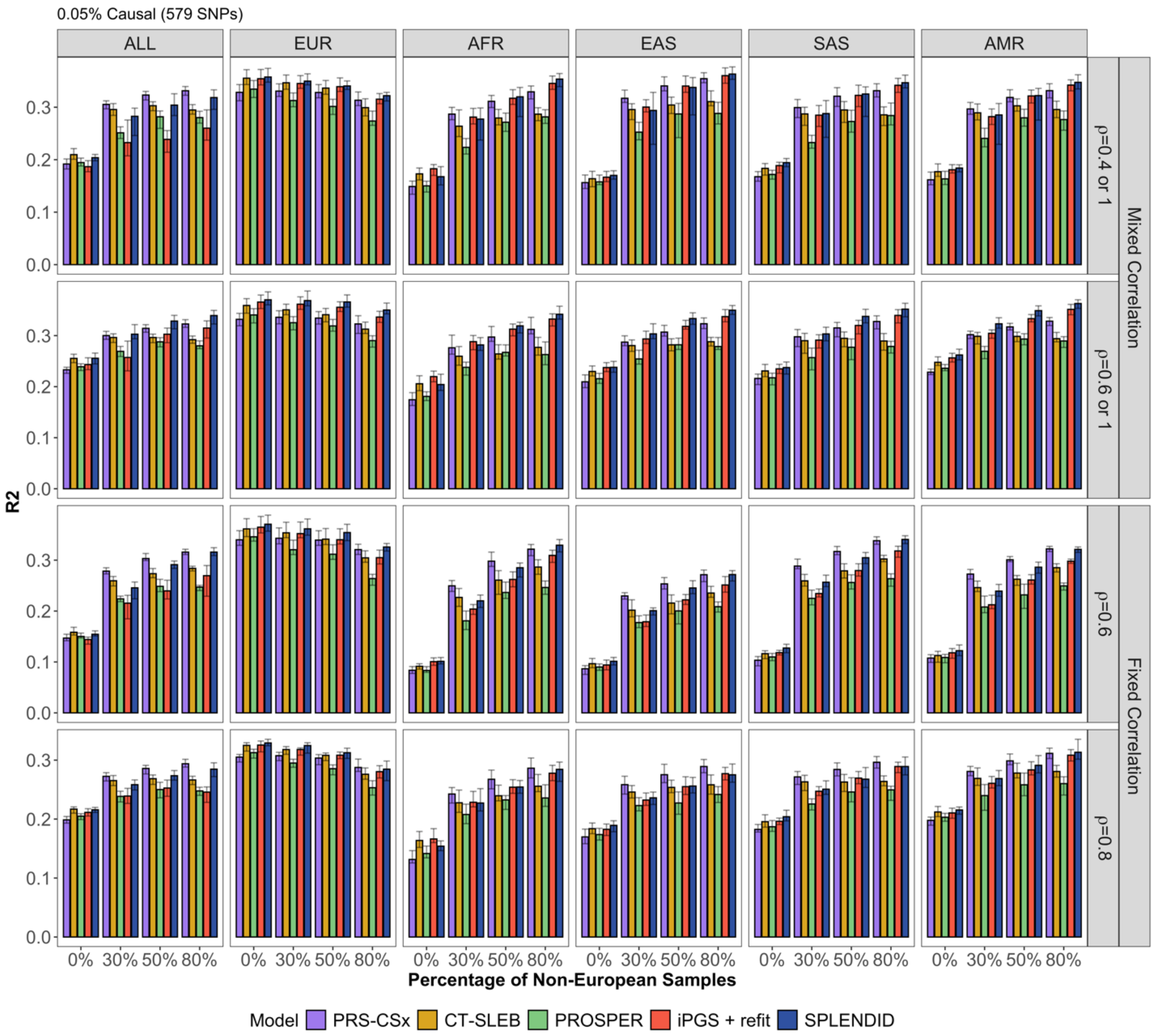


**Supplementary Figure 1 Prediction for 0.05% causal proportion**. We compare SPLENDID against existing methods – PRS-CSx, CT-SLEB, PROSPER, and iPGS+refit – with varying ancestry proportions and genetic correlation patterns. In this highly sparse setting, SPLENDID outperforms all methods when heterogeneity is mixed (top two panels), and performs similar to PRS-CSx under fixed genetic correlation (bottom two panels).

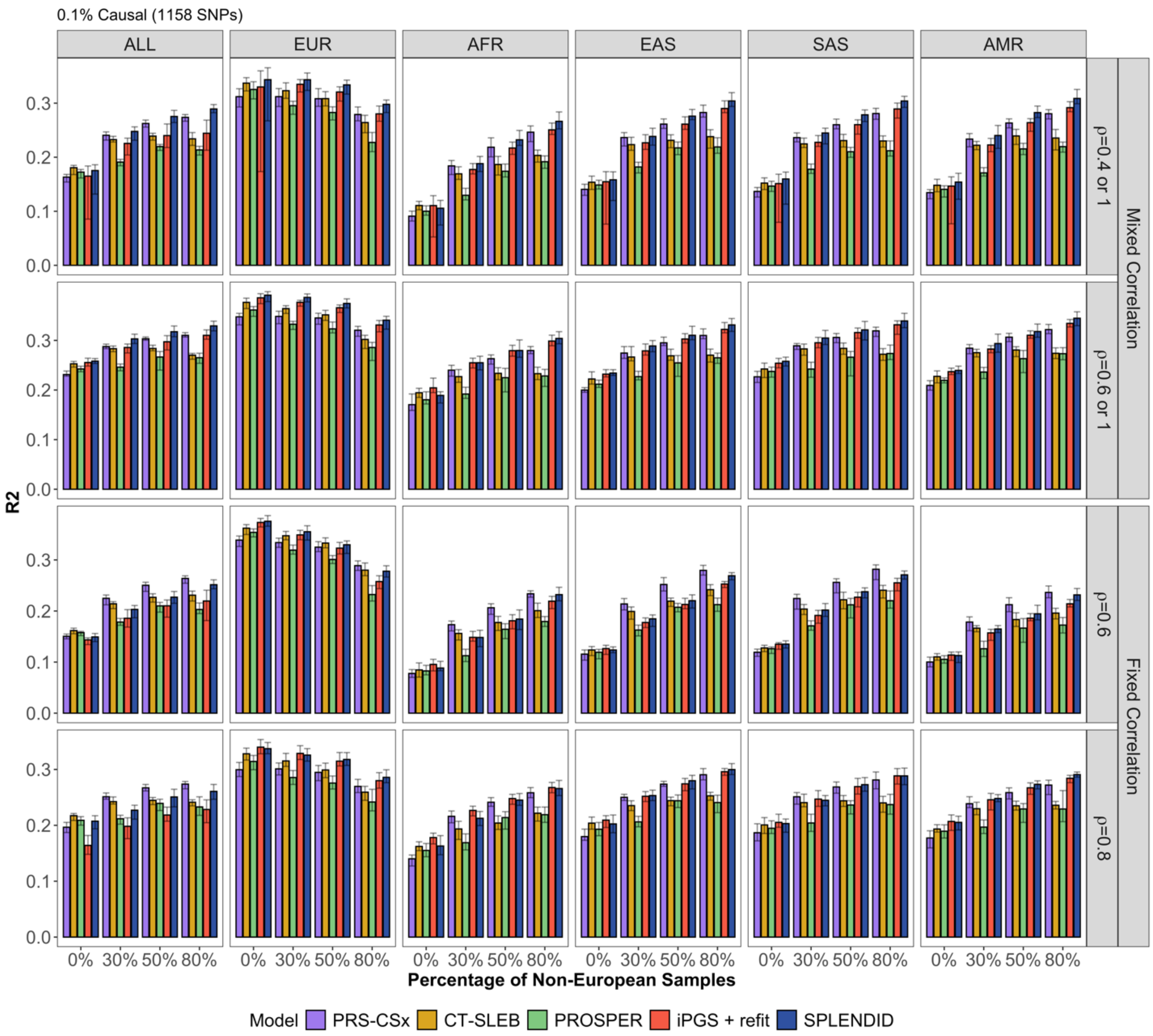


**Supplementary Figure 2 Prediction for 0.1% causal proportion.** We compare SPLENDID against existing methods – PRS-CSx, CT-SLEB, PROSPER, and iPGS+refit – with varying ancestry proportions and genetic correlation patterns. In this relatively sparse setting, SPLENDID outperforms all methods when heterogeneity is mixed (bottom two panels), and performs similar to PRS-CSx for a constant genetic correlation across ancestry (bottom two panels).

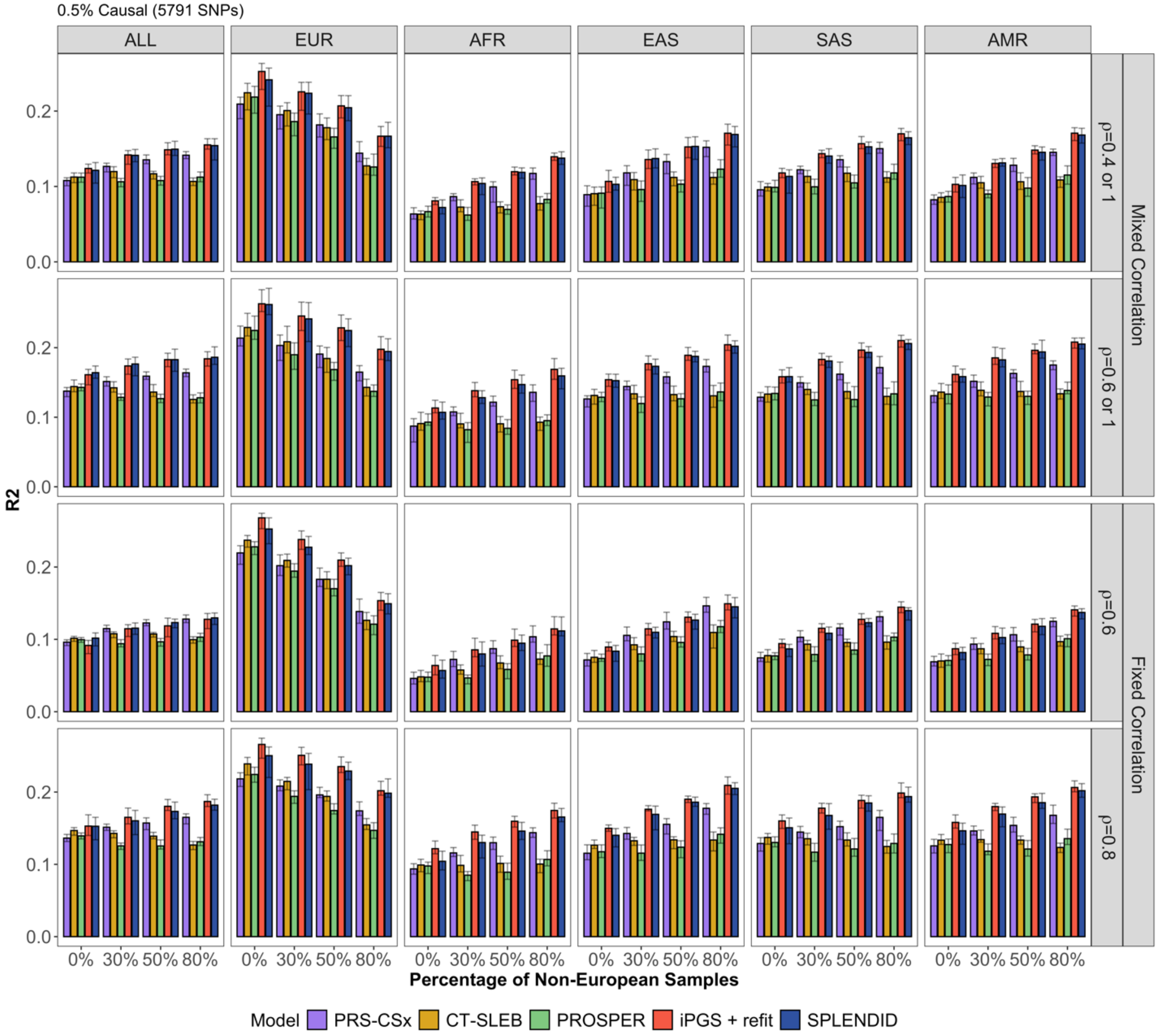


**Supplementary Figure 3. Prediction for 0.5% causal proportion.** We compare SPLENDID against existing methods – PRS-CSx, CT-SLEB, PROSPER, and iPGS+refit – with varying ancestry proportions and genetic correlation patterns. In this relatively dense setting, SPLENDID outperforms all GWAS-based methods, and performs similarly or slightly below iPGS+refit.

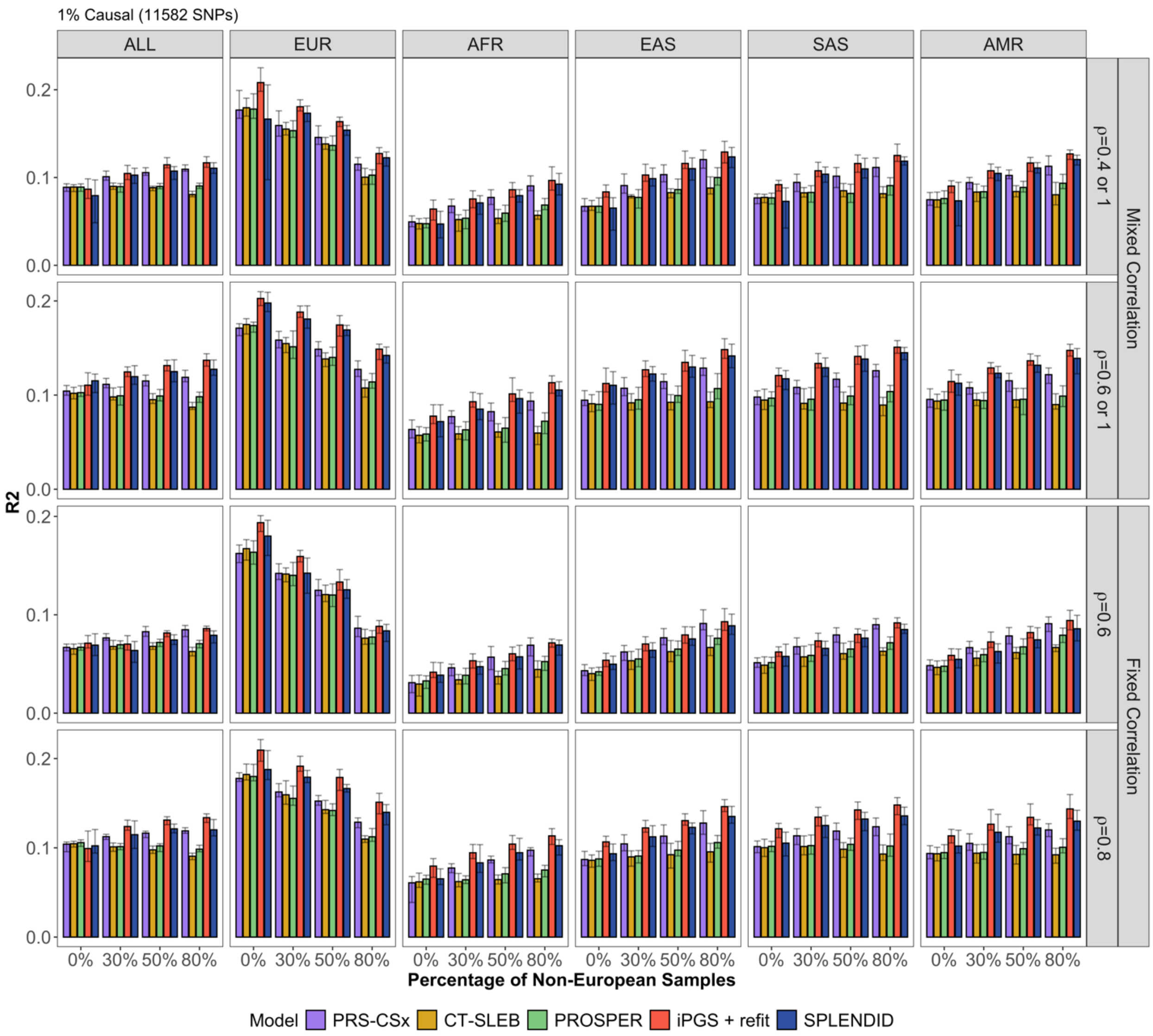


**Supplementary Figure 4. Prediction for 1% causal proportion.** We compare SPLENDID against existing methods – PRS-CSx, CT-SLEB, PROSPER, and iPGS+refit – with varying ancestry proportions and genetic correlation patterns. In this relatively dense setting, SPLENDID outperforms all GWAS-based methods, and performs similarly or slightly below iPGS+refit.

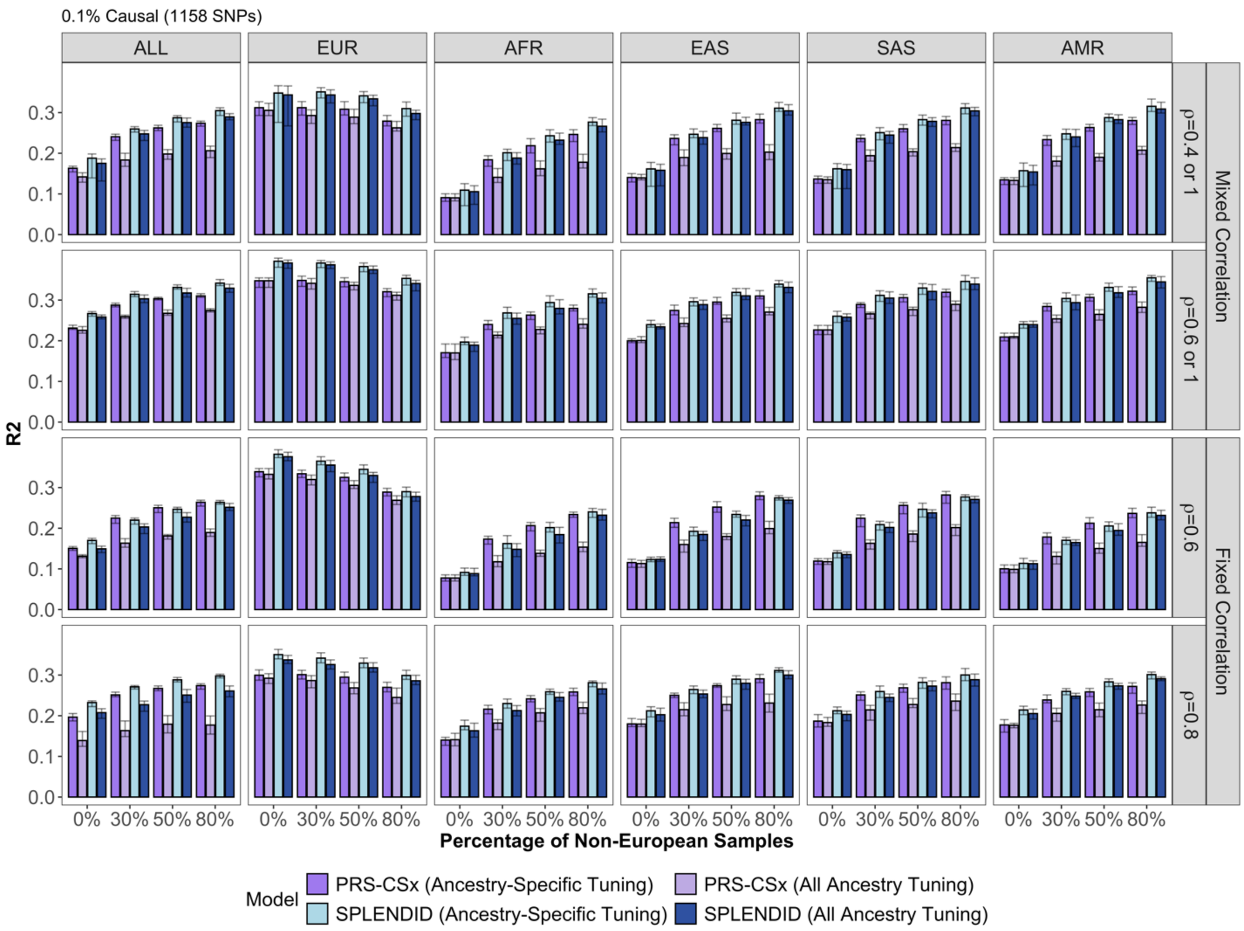


**Supplementary Figure 5. Comparing ancestry-specific and all-ancestry tuning for PRS-CSx and SPLENDID under 0.1% causal proportion.** PRS-CSx performs similar to SPLENDID in sparse settings and substantially better than similar GWAS-based approaches. We plot the prediction accuracy of PRS-CSx and SPLENDID when tuned within each ancestry – as designed for PRS-CSx – and pooled across all ancestries – as designed for SPLENDID. SPLENDID performed similarly in both cases, while PRS-CSx only performs well with ancestry-specific tuning. This shows the robustness of SPLENDID to tuning data, particularly for creating a single PRS model across several ancestries.

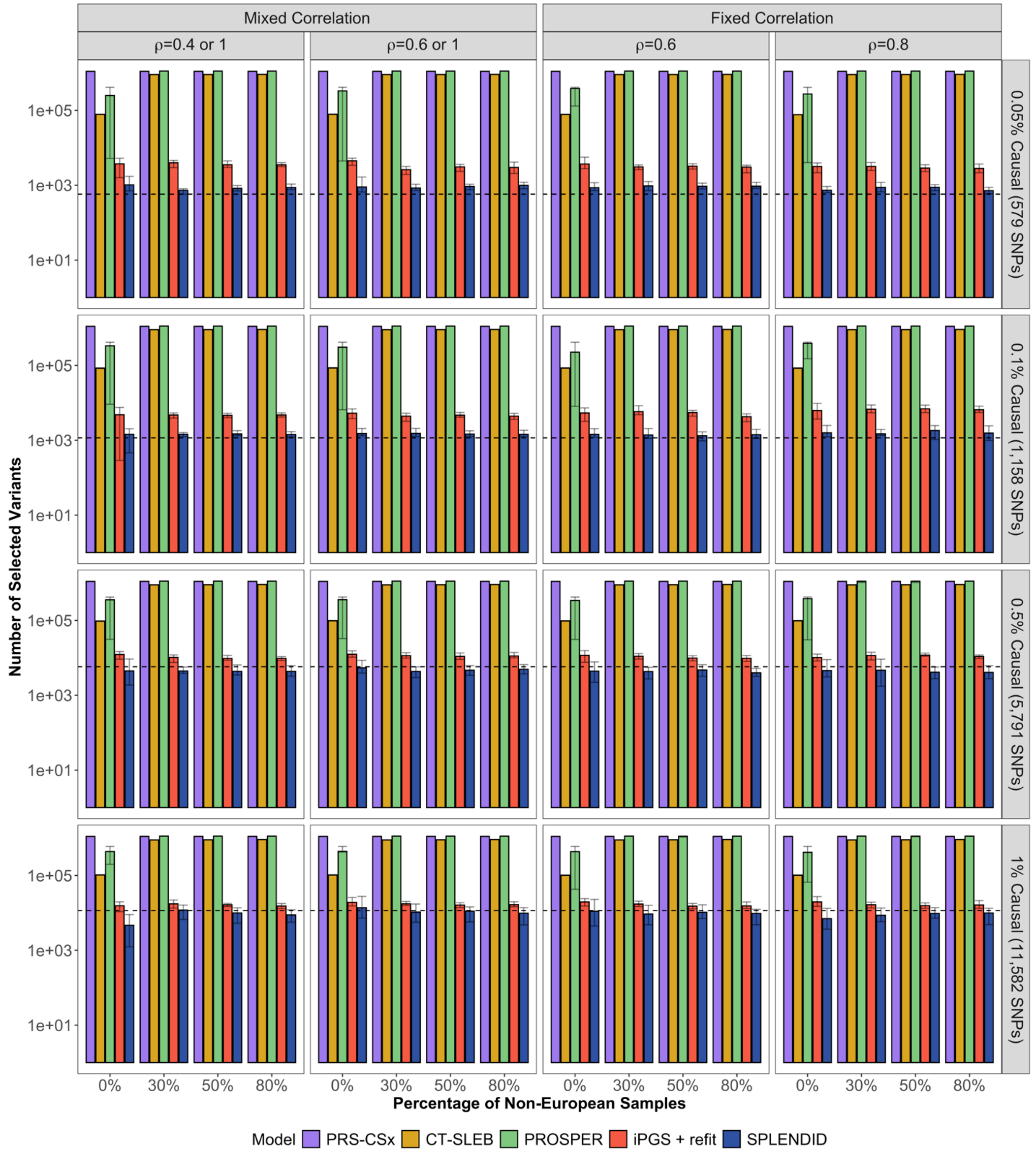


**Supplementary Figure 6 Variable selection for five PRS methods in simulation studies.** Across both sparse and dense settings, SPLENDID selected the fewest number of variants in the PRS and is similar to the true number of causal variants (dashed line). Most GWAS-based approaches select a very large number of variants, while Lasso in iPGS+refit also tends to over-select compared to an L0L1-based penalty in SPLENDID.

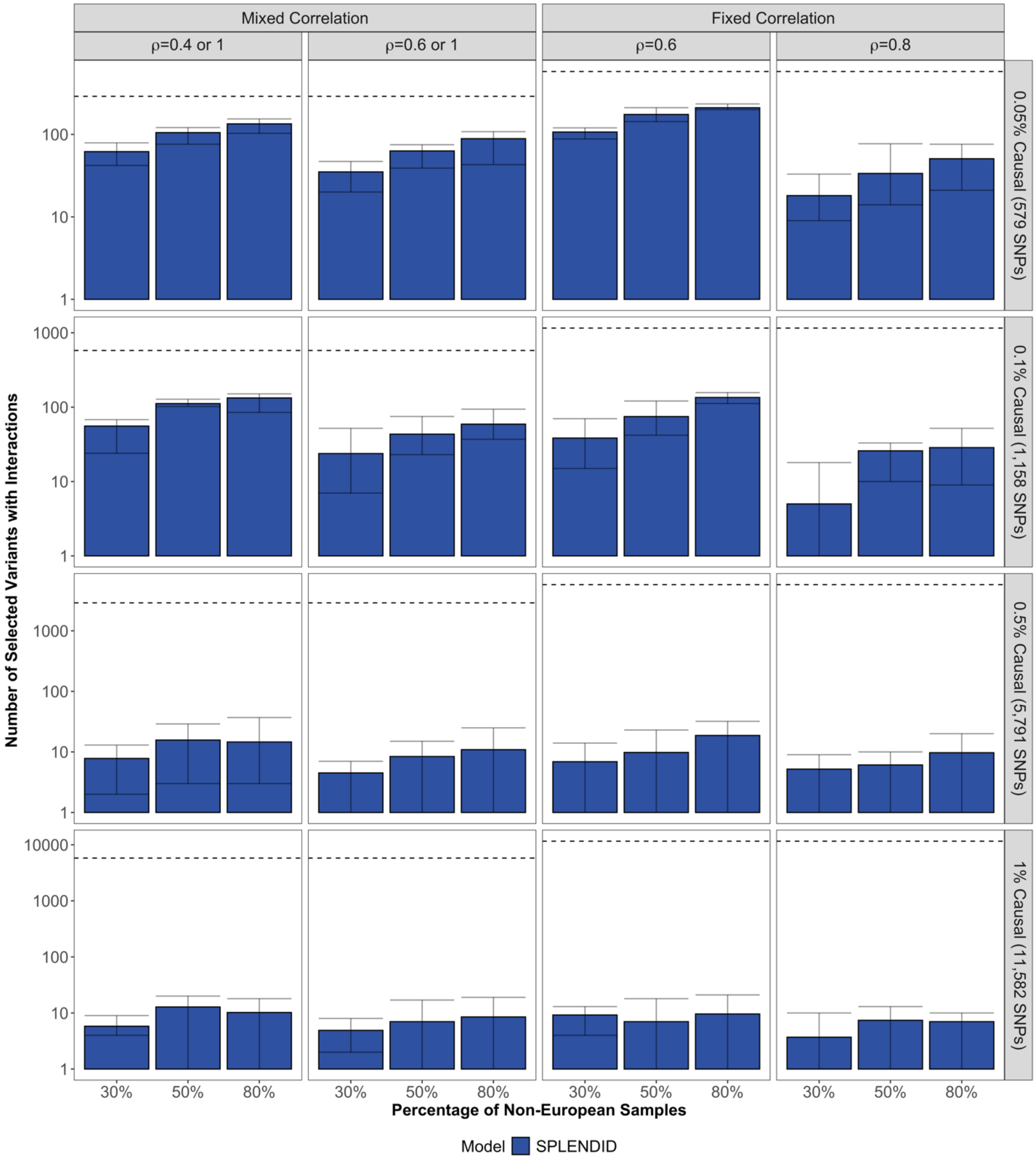


**Supplementary Figure 7 Number of selected interactions by SPLENDID in simulation studies.** We plot the average number of variants (on the log-10 scale) selected with interactions by SPLENDID across different causal variant proportions, ancestry proportions, and genetic correlation patterns. Dashed horizontal lines indicate the number of variants with heterogeneous effects (i.e. “true” number of interactions). In the sparse settings (top two rows), higher non-European ancestry proportion leads to selection of more interactions. Meanwhile, in denser settings (bottom two rows), only a small number of interactions are selected at any ancestry proportion level due to the relatively small magnitude of heterogeneity in these settings.

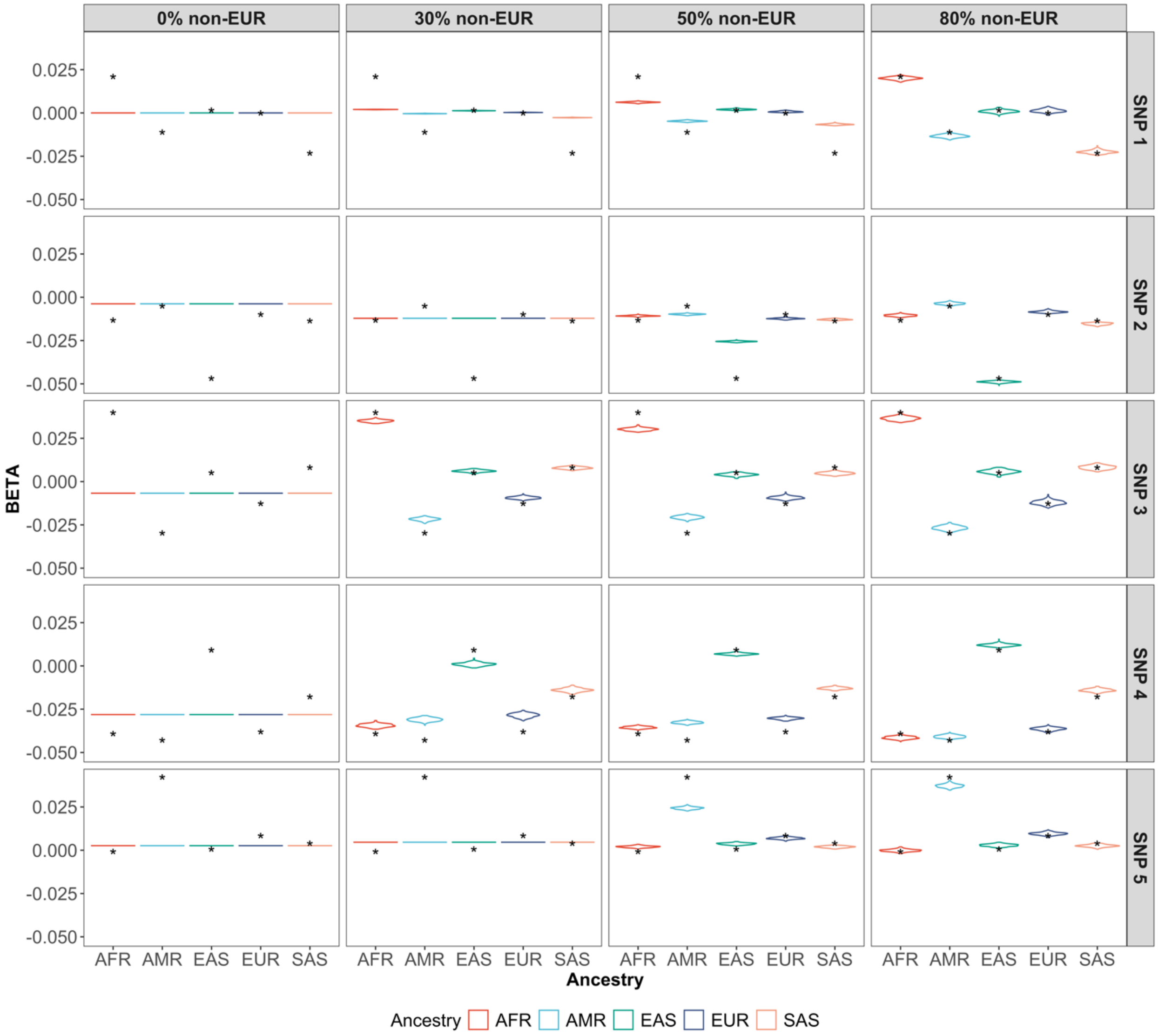


**Supplementary Figure 8 Example of estimated individual-level effect sizes for 0.1% causal variants and 60% fixed genetic correlation.** We plot individual-level effect sizes, according to GxPC interactions, for 5 causal variants correctly selected by SPLENDID, averaged over 10 simulation replicates. As the proportion of non-European training data increases (top panels), the estimated effect sizes (colored violin plots) become much closer to the true effect sizes for each ancestry (stars).

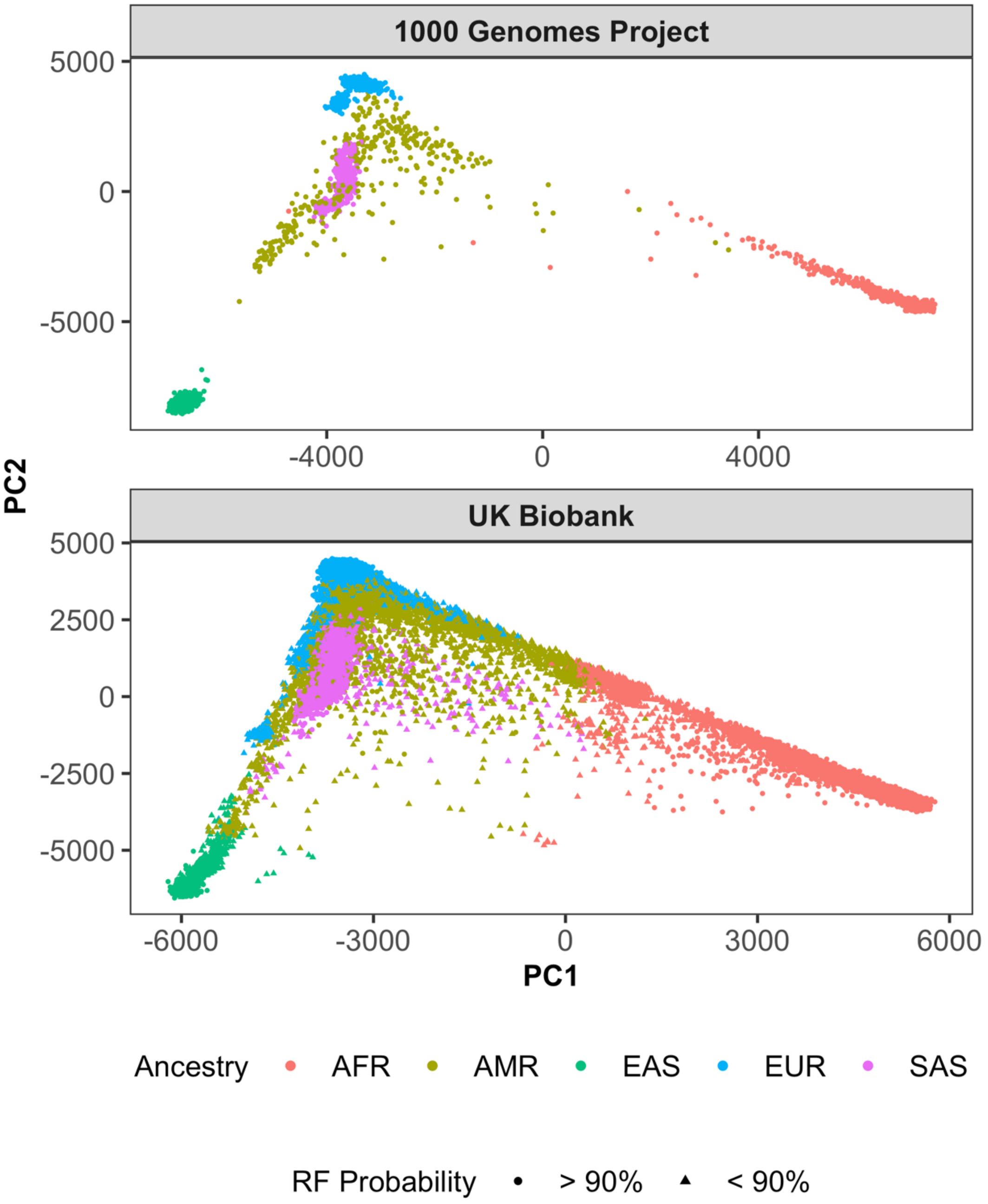


**Supplementary Figure 9 Ancestry principal components analysis across 1000 Genomes Project Phase 3 and UK Biobank.** We conducted PCA within 1000 Genomes Project Phase 3 (1000G), and projected genotypes from UK Biobank (UKBB) onto this PC space. We show the distribution of genetic ancestry in each dataset based on the first two ancestry PCs. For 1000G, samples are colored by pre-defined population labels. For UKBB, we first trained a random forest classifier using the first 20

PCs in 1000G according to population labels, and then applied this to the projected PCs in UKBB to generate predicted ancestry labels. Triangles in the UKBB panel denote samples for which the random forest probability of the assigned ancestry is below 90%.

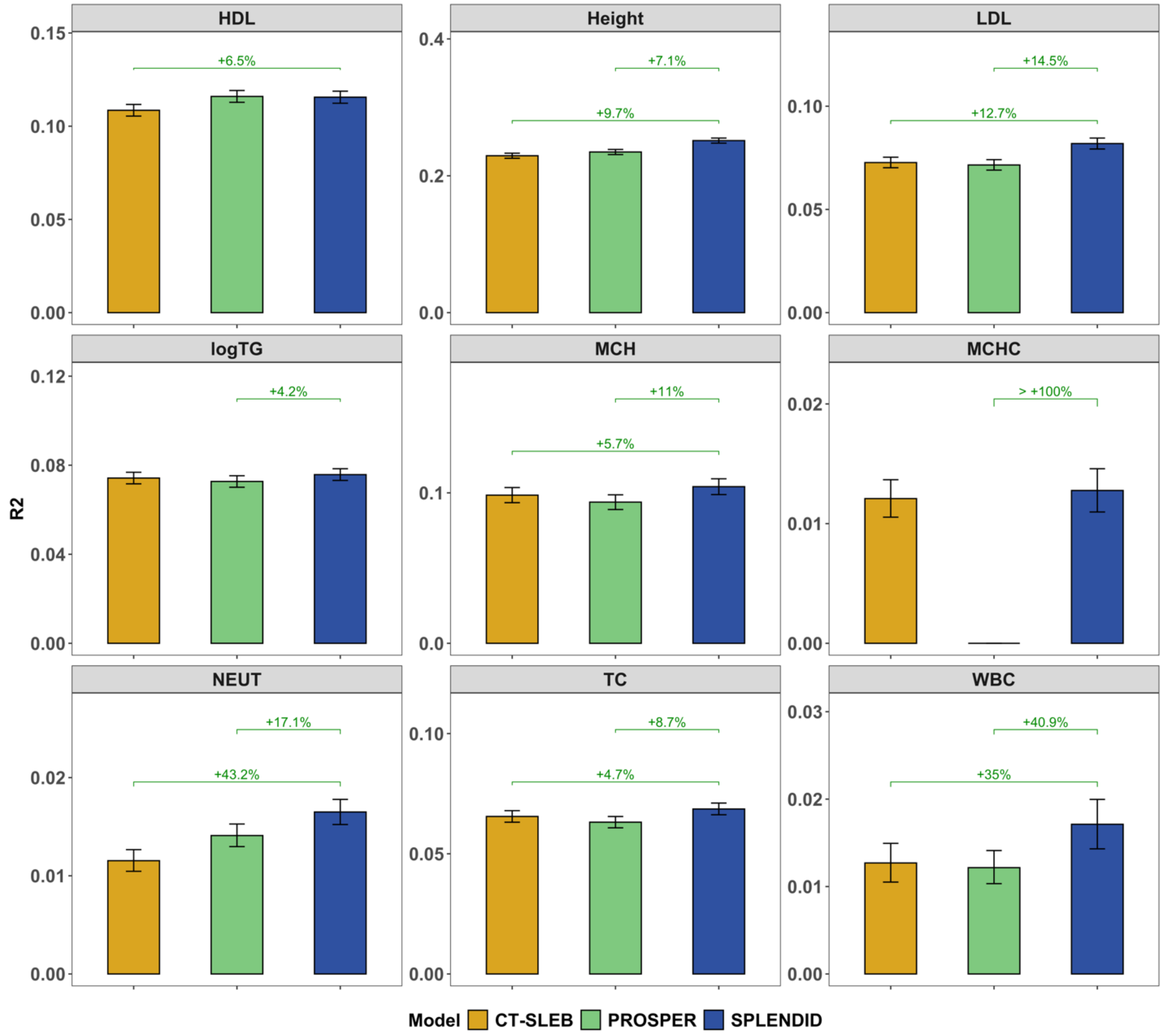


**Supplementary Figure 10 Prediction accuracy of CT-SLEB, PROSPER, and SPLENDID for EUR with reduced EUR sample size in UKBB tuning data.** In our main analyses, CT-SLEB and PROPSER often achieved higher $R^2$ than SPLENDID in EUR ancestry, which could be attributed to their ancestry-specific SuperLearner steps that take advantage of the large EUR sample size in UKBB. Thus, we compared prediction accuracy of SPLENDID versus CT-SLEB and PROPSER in EUR after reducing the EUR sample size in UKBB tuning from about 160,000 to 50,000, where neither CT-SLEB nor PROSPER outperformed SPLENDID. Adjusted $R^2$ was computed in the UKBB validation dataset, with error bars corresponding to the 95% BCI for adjusted $R^2$ using 10,000 replicates. 95% BCI were also constructed for the percent increase in $R^2$ by using external GWAS meta-analysis, where values are shown for results where this 95% BCI did not include 0.

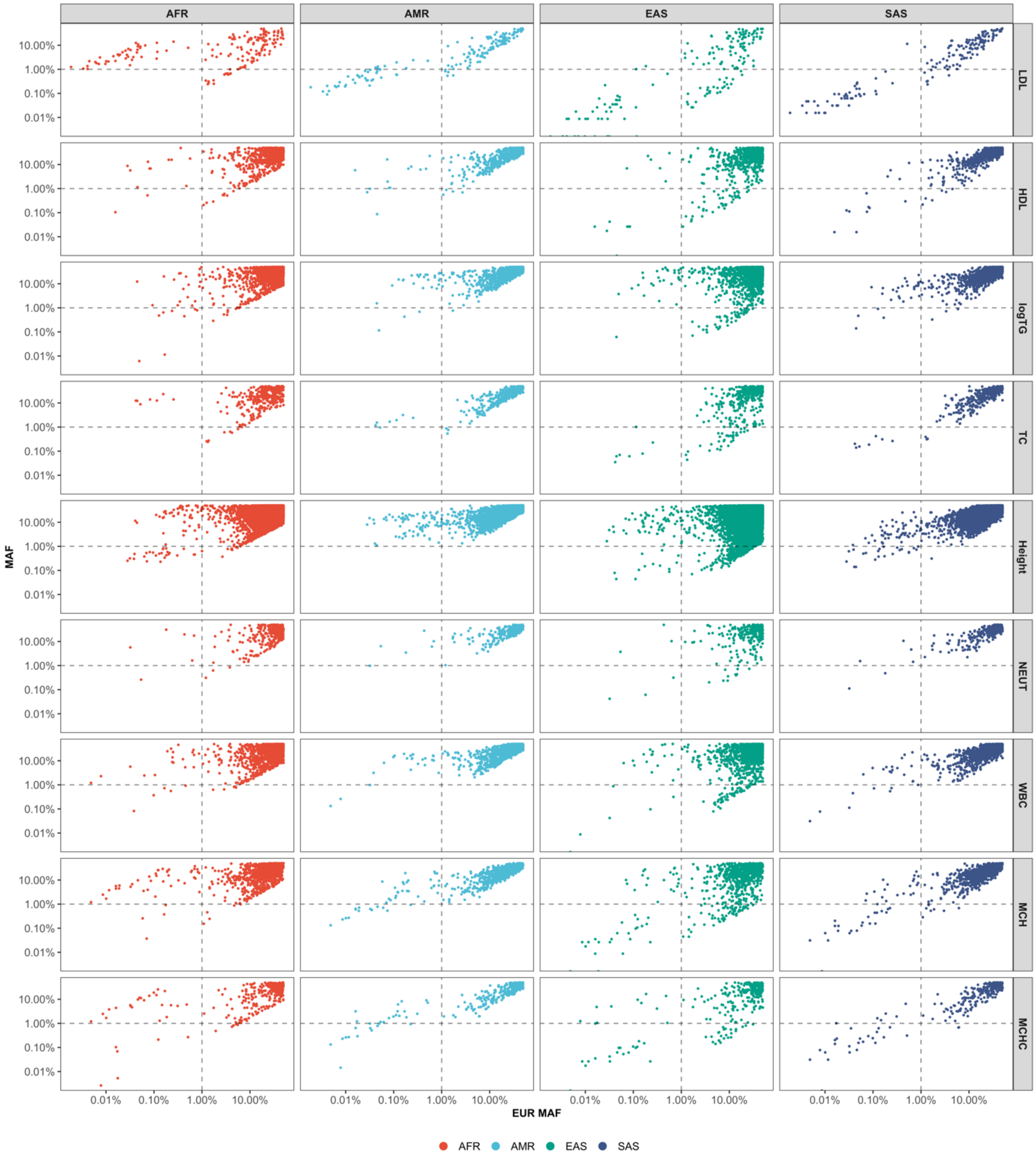


**Supplementary Figure 11 Allele frequency of selected variants from SPLENDID in different ancestries within All of Us**. We compare allele frequencies of variants selected by SPLENDID for each of the 9 continuous traits in All of Us Research Program, which varies immensely across different ancestries. The X-axis shows the MAF for European ancestry in AoU, while the Y-axis shows the MAF for each of the four non-European ancestries (AFR, AMR, EAS, SAS). Dotted lines indicate MAF of 1% to separate common variants from rare and low-frequency variants in each ancestry.

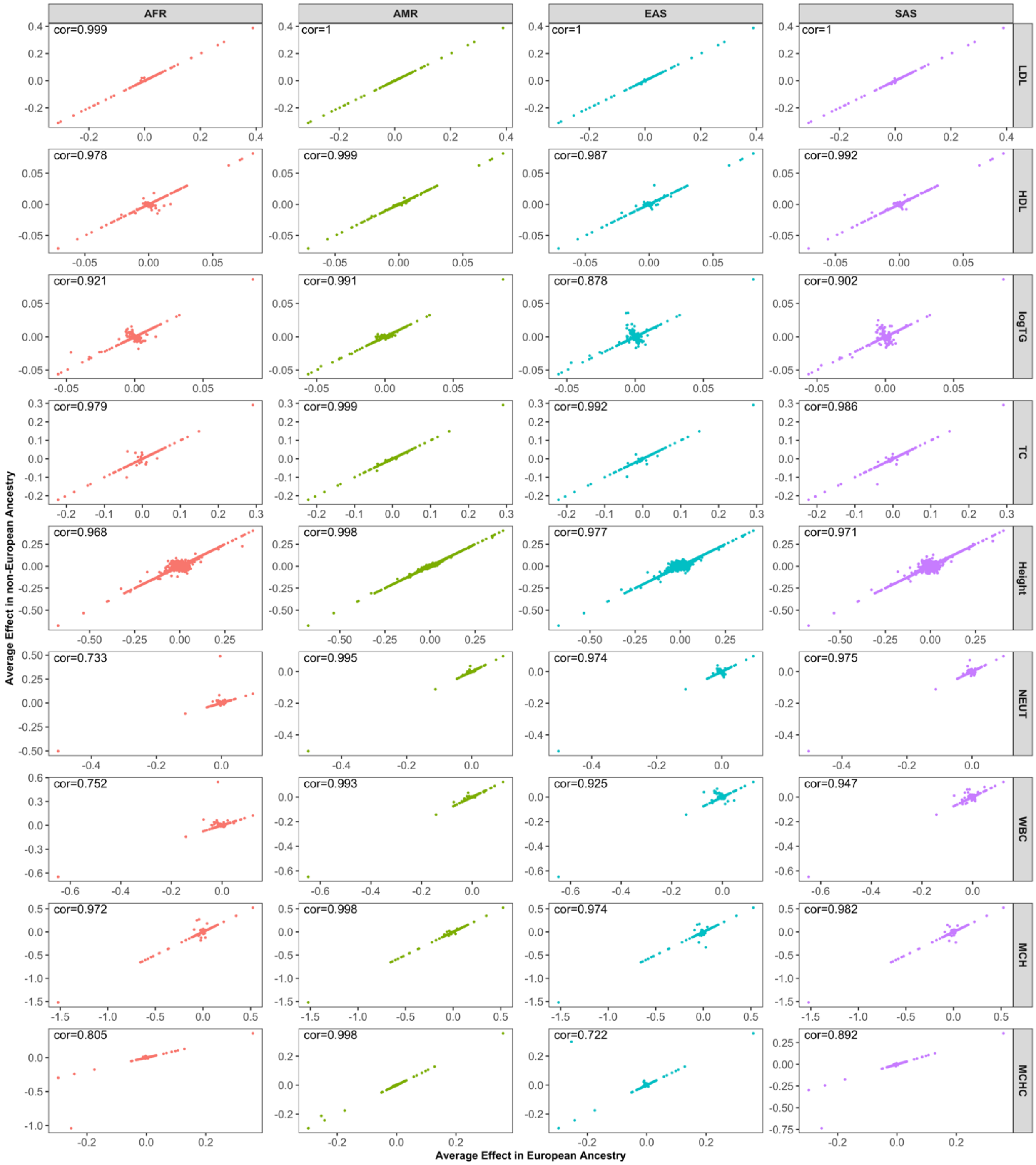


**Supplementary Figure 12 Estimated effect sizes by ancestry in UK Biobank validation data.** We compared estimated effect sizes, incorporating GxPC interactions, for variant selected by SPLENDID for each of the 9 continuous traits in All of Us Research Program. X-axis shows the average effect size of each variant for EUR, and the Y-axis shows the average effect size in non-European ancestry. Results are stratified by trait (side panels) and genetic ancestry (top panels), with text showing the correlation in effect estimates between European and non-European ancestries.

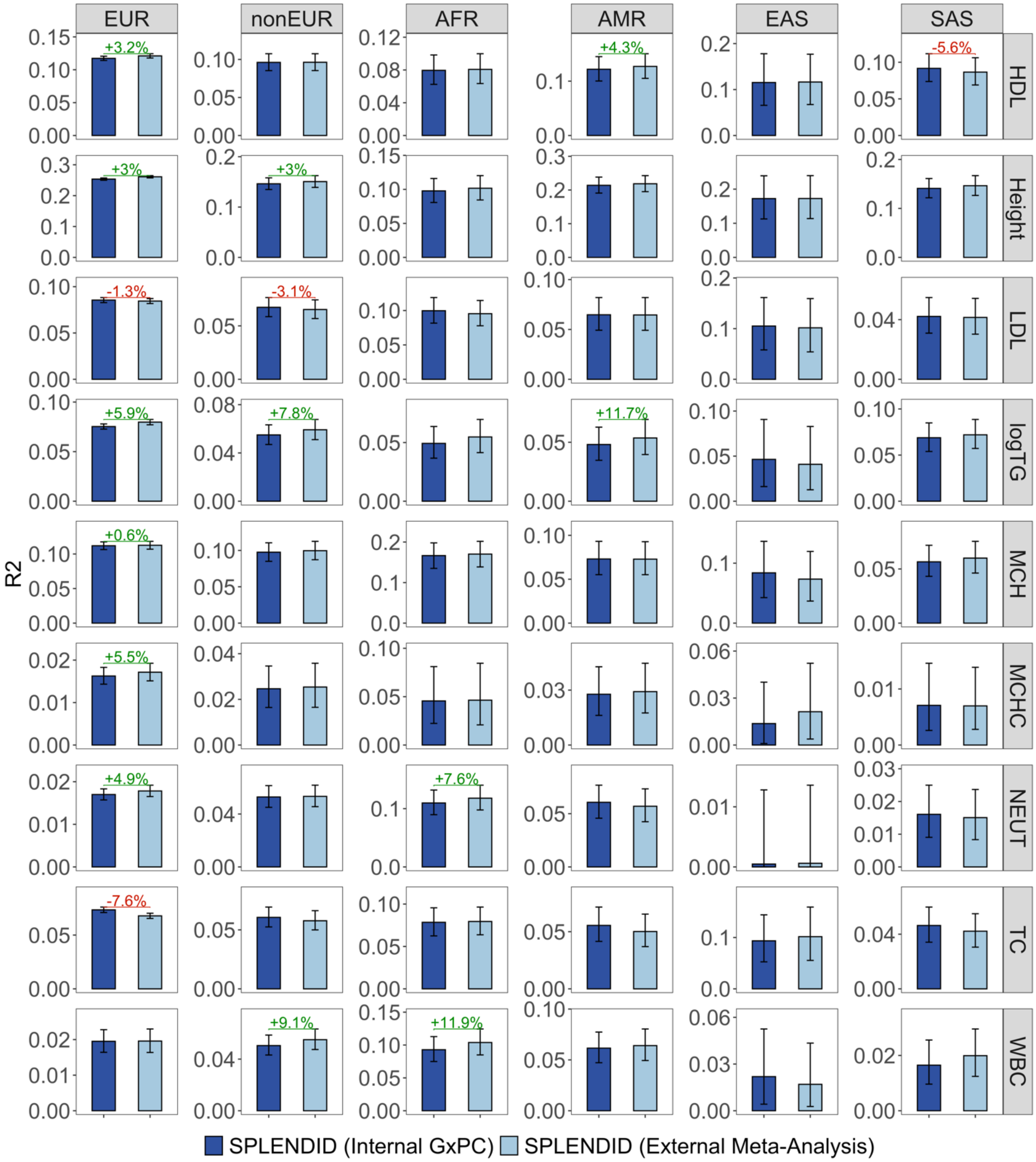


**Supplementary Figure 13. Comparing prediction accuracy of SPLENDID using GxPC interaction GWAS in All of Us or meta-analysis of published ancestry-specific GWAS.** We compared prediction accuracy of SPLENDID when using p-values from GxPC interaction GWAS in AoU training data or meta-analysis of external GWAS to select interactions in the regression model. Adjusted $R^2$ was computed in the UKBB validation dataset, with error bars corresponding to the 95% BCI for adjusted $R^2$ using 10,000 replicates. 95% BCI were also constructed for the percent increase in $R^2$ by using external GWAS meta-analysis, where values are shown for results where this 95% BCI did not include 0.

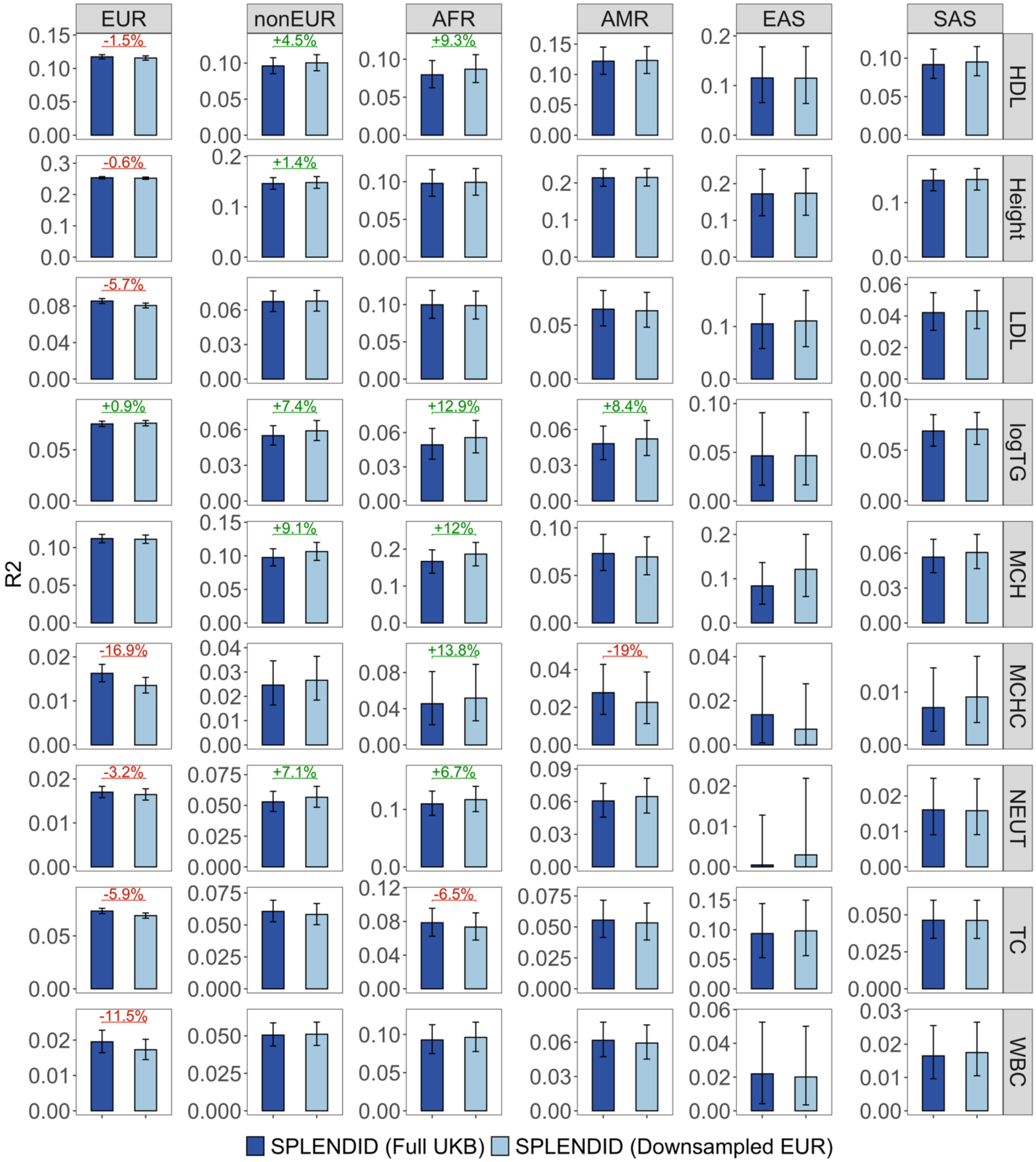


**Supplementary Figure 14. Comparing prediction accuracy of SPLENDID using full UK Biobank and down-sampled EUR.** We evaluated the robustness of SPLENDID to the ancestry proportions for PRS tuning by comparing between using the full UKBB tuning dataset (about 93% EUR) and a down-sampled tuning dataset with about 30% EUR. Adjusted $R^2$ was computed in the UKBB validation dataset, with error bars corresponding to the 95% BCI for adjusted $R^2$ using 10,000 replicates. 95% BCI were also constructed for the percent increase in $R^2$ by using the down-sampled tuning dataset, where values are shown for results where this 95% BCI did not include 0.

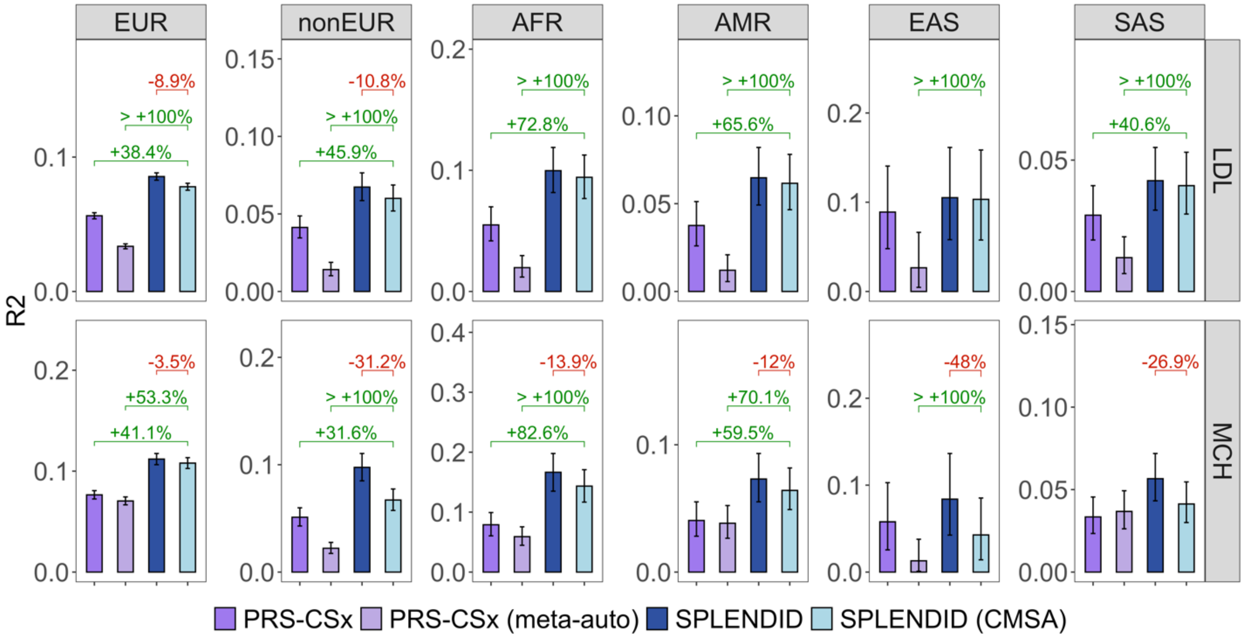


**Supplementary Figure 15. Comparing prediction of LDL and MCH by SPLENDID and PRS-CSx in UKBB with and without tuning data**. We compared the performance of implementations for SPLENDID and PRS-CSx with and without an independent tuning dataset. For SPLENDID, we adopted a CMSA approach that averaged results across different folds within the AoU tuning data. We compared this with PRS-CSx-meta-auto, which automatically selects prior hyperparameters and combines posterior effect estimates from each ancestry, and the original implementations of SPLENDID and PRS-CSx that used the UKBB tuning dataset. Adjusted $R^2$ was computed in the same UKBB validation dataset, with error bars corresponding to the 95% BCI for adjusted $R^2$ using 10,000 replicates. 95% BCI were also constructed for the percent increase in $R^2$ by the CMSA version of SPLENDID, where values are shown for results where this 95% BCI did not include 0.
